# Metagenomic-based network analysis reveals the importance of vitamin cross-feeding in marine microbial assemblages

**DOI:** 10.1101/2025.08.08.668683

**Authors:** Nestor Arandia-Gorostidi, Lidia Montiel, Francisco Latorre, Vanessa Balagué, Josep M. Gasol, Rafel Simó, Caterina R. Giner, Ramiro Logares

## Abstract

Vitamins play a fundamental role in microbial metabolism and interactions, yet their scarcity in marine environments and the limited ability for de novo synthesis of some vitamins often leads to metabolic dependencies and cross-feeding among microbial taxa. Using a decade-long time-series metagenomics dataset from the Blanes Bay Microbial Observatory (BBMO), we investigated the role of vitamin biosynthesis in structuring the seasonal marine microbial interactions among prokaryotes and eukaryotes. Gene co-occurrence analysis revealed that vitamin-related metabolism had the highest number of associations among all metabolic pathways, underscoring the potential role of vitamins in microbial interactions. Metagenome-assembled genome (MAG) co-occurrence analysis further identified the prokaryotic biosynthesis of cobalamin (B_12_) and thiamine (B_1_) as key mediators of these associations. Rather than between complete vitamin synthesizers and auxotrophs, the main associations were found between partial synthesizers, suggesting that vitamer cross-feeding may represent an essential microbial interaction in sustaining vitamin biosynthesis throughout the year. While complete cobalamin synthesizers dominate in summer, partial synthesizers and lower-ligand activators are more prevalent in winter, driving cross-feeding interactions. In contrast, thiamine biosynthesis is more widespread, with complete synthesizers peaking in winter. The association strength, however, was maximum between prokaryotic vitamin producers and eukaryotes, highlighting the dependence of eukaryotes on bacterial vitamin production. These results provide novel insights into seasonal metabolic interdependencies, emphasizing the ecological significance of vitamin biosynthesis in marine ecosystems.

## INTRODUCTION

Interactions among marine microorganisms largely shape the structure and dynamics of marine microbiomes. These interactions are essential across key biogeochemical processes, from the nitrogen cycle, where associations between nitrogen fixers, ammonia oxidizers, and nitrate reducers sustain oceanic nitrogen cycling (Zehr & Kudela, 2011; Pachiadaki et al., 2017), to carbon cycling, in which the association between diverse microbial taxa control the degradation and remineralization of organic carbon (Datta et al., 2016). However, understanding how microbes interact, either through cooperation or competition, in natural environments remains a significant challenge due to the complexity and diversity of microbial communities, as well as the difficulty of directly observing their interactions *in situ*. As a result, the mechanisms and ecological implications of microbial cooperation remain poorly understood.

Studies utilizing co-cultures have started to uncover the mechanisms driving specific interactions between marine microorganisms (Hibbing et al., 2010). These interactions are driven by the release and subsequent uptake of a myriad of metabolites produced as a result of microbial metabolism, such as vitamins, amino acids, organic acids, or signaling molecules. The release of these secondary metabolites can profoundly influence the growth and behavior of community members, playing a crucial role in shaping the metabolism and assembly of the entire marine microbial community (Durham et al., 2015; Oña & Kost, 2022; Moran et al., 2022; Durham, 2021). Among these compounds, vitamins have garnered significant attention in recent studies due to their essential role across nearly all forms of life (Croft et al., 2006; Giovannoni, 2012; Combs & McClung, 2017), despite their widespread scarcity in marine environments (Panzeca et al., 2009; Sañudo-Wilhelmy et al., 2012). In particular, B_1_ (Thiamine) and B_12_ (Cobalamin) are the most required vitamins among microbes (Sañudo-Wilhelmy et al., 2014), serving as main cofactors in central carbon metabolism (Berg et al., 2007; Dowling et al., 2012). In marine ecosystems, microbial biosynthesis is the primary source of both B_1_ and B_12_. Yet, due to the genetic and metabolic costs associated with these biosynthetic pathways, many microorganisms lack complete de novo synthesis capabilities. This limitation not only promotes mutualistic relationships between vitamin synthesizers and auxotrophs (Croft et al., 2005; Suffridge et al., 2018) but also leads to more complex interactions where different microbes contribute to the full synthesis of a given vitamin through the synthesis of vitamin precursors or vitamers (Suffridge et al., 2020; Paerl et al., 2023). However, the limited number of studies addressing interactions related to vitamin synthesis within natural microbial assemblages constrains our understanding of their importance in mediating microbial interactions and nutrient dependencies in marine environments.

Although co-culture studies have provided valuable insights into microbial interactions (Sultana et al., 2023; Wienhausen et al., 2024), their scope remains limited due to the challenges of cultivating many marine microorganisms. Computational tools have become increasingly important over the past two decades for generating hypotheses about microbial interactions in natural environments. For instance, taxonomic co-occurrence and co-exclusion analyses over space or time have provided valuable insights into potential interactions among marine microorganisms, including bacteria-bacteria (Lima-Méndez et al., 2015; Deutschmann et al., 2024), and bacteria-phytoplankton interactions (Arandia-Gorostidi et al., 2022), suggesting that microbial interactions influence the seasonal community structure. Even though taxonomic association analyses (based on co-occurrences and co-exclusions) help to describe hypothetical ecological interactions, they provide very limited information on the mechanisms underlying these interactions. Gene-level co-occurrence analyses through metagenomics, on the other hand, offer a more granular view of the potential metabolic bases of microbial interactions (Fondi et al., 2016; Fondi & Patti, 2019), pointing out key metabolites as drivers of these metabolic associations. Recent computational models based on metagenomics have proven highly valuable in uncovering microbial interactions across diverse environments, including hydrogen sulfide-related interactions among hydrothermal microbes (Kuppa Baskaran et al., 2023) and associations mediated by amino acid and vitamin prototrophy and auxotrophy in the global ocean (Giordano et al., 2024).

In this study, we aimed to investigate the metabolic bases of the most abundant marine microbial interactions by analyzing a 12-year metagenomic time-series dataset from the Mediterranean Sea. By combining the gene databases with co-occurrence analyses, we examine the key metabolic pathways and potential metabolites that underpin microbial interactions within the Bay of Blanes community over a decade. Furthermore, we leverage metagenome-assembled genomes (MAGs) to identify potential interactions between prokaryotic groups and characterize their seasonal vitamin-related interactions.

## MATERIAL & METHODS

### Sample collection and DNA extraction

Monthly seawater samples for metagenomic analyses were collected at the Blanes Bay Microbial Observatory (BBMO; http://bbmo.icm.csic.es) in the Northwestern Mediterranean Sea (41°40’N, 2°48’E). Sampling was conducted at the surface (≈ 1 m depth) between January 2009 and December 2020. Approximately 6 L of seawater was prefiltered first through a 200 μm mesh, and subsequently through a 3 μm pore-size polycarbonate filter (Poretics), and a 0.2 μm Sterivex Millipore filter using a peristaltic pump. For the metagenomics analysis, only the fraction between 3 μm and 0.2 μm was used. The Sterivex units were stored at −80 °C until further analysis. Besides metagenomic analyses, samples were collected to measure total chlorophyll-a concentration (*µ*g/l), inorganic nutrients (PO_4_^3*−*^, NH_4_^+^, NO_2_*^−^*, NO_3_*^−^*, and SiO_2_), temperature, and salinity. Water temperature and salinity were sampled in situ using an SAIV A/S SD204 CTD. Chlorophyll *a* (Chl *a*) concentrations were analyzed by fluorometry (10-AU Fluorometer Turned Designs). Inorganic nutrients were measured using an Alliance Evolution II autoanalyzer (Graßhoff et al., 2009). Cell counts to determine bacterial and picoeukaryote total abundance were done by flow cytometry as described in Gasol et al. (2016).

Once in the laboratory, DNA was extracted with a standard phenol–chloroform protocol as described in Massana et al. (1997) and purified using an Amicon unit (Millipore). Sample purification was performed as described in Krabberød et al. (2022). Shotgun sequencing of the samples was performed in separate batches. Samples between 2009 and 2012 were sequenced using an Illumina Hiseq4000 (2 x 150 bp), while the samples between 2013 and 2020 were sequenced using the Illumina NovaSeq6000 (2 x 150 bp) platform. All samples had a sequencing coverage > 30 gigabases (Gb).

### Trimming, assembly, and gene prediction

Adapters and low-quality reads were removed from the samples using Cutadapt (v3.5, Martin, 2011). Each sample was subsequently assembled individually with Megahit (v1.1.3) to generate contigs (Li et al., 2015) using the meta-large option. Protein-coding regions in the contigs were predicted for each sample using Prodigal (v2.6.3, Hyatt et al., 2010) and MetaGeneMark (v3.38, Gemayel et al., 2022), considering partial Open Reading Frames (ORFs) and a minimum fragment length of 250 bp. The resulting dataset, containing 500 million genes, was clustered at 95% identity and 80% coverage of the shorter sequence using Linclust v10 (Steinegger & Söding, 2018), resulting in a final catalog of 231 million genes (ORFs). Predicted genes were functionally annotated with the Kyoto Encyclopedia of Genes and Genomes (KEGG, Kanehisa & Goto, 2000), using Diamond blastp (Buchfink et al., 2015). Genes were taxonomically annotated against the Genome Taxonomy Database (GTDB) release 95 (Parks et al., 2018) using MMSeqs2 (v9-d36de, Steinegger et al., 2017) with a sensitivity value of 5.7, specifying ranks at which the Lowest Common Ancestor (LCA) should be determined.

Gene abundances per sample were calculated by mapping metagenome reads back to the catalogue using BWA (v0.7.17, Li et al., 2009) and obtaining the number of counts per gene using HTSeq (v0.10.0, Anders et al., 2015). Gene counts were normalized within and among samples by gene length and the abundance of 10 single-copy genes as done by others (Sunagawa et al., 2013; Milanese et al., 2019). The generated normalized gene abundance tables include the abundance of each gene (ORF) in each sample. From these, we calculated the corresponding functional abundance tables by adding all the normalized abundances of all genes annotated to a specific KEGG function.

### Metagenome-assembled genomes (MAG)

To reconstruct metagenome-assembled genomes (MAGs), a total of 84 metagenomes from the Blanes Bay Microbial Observatory (BBMO) were used. We used Simka (v1.5, Benoit et al., 2016) to estimate Bray-Curtis dissimilarities among metagenomes using 21-mer profiles. These dissimilarities were then clustered using hierarchical clustering (hclust) in R, allowing us to define four distinct metagenomic groups (G1 to G4), which also corresponded to seasonal groupings: G1 (Winter), G2 (Spring), G3 (Early Summer), and G4 (Late Summer). Two metagenomes were excluded during this step as they did not cluster with any group, resulting in 82 metagenomes in total.

Each group of metagenomes was co-assembled separately using MEGAHIT (v1.2.8., Li et al., 2015). Reads from the original metagenomes were then mapped back to their respective co-assemblies using BWA (v0.7.17-r1188, Li et al., 2009) and SAMtools (v1.8 for G2, G3, G4 and v1.12 for G1, Danecek et al., 2021). MAGs were first generated using MetaBAT 2 (v2.12.1, Kang et al., 2019) with default parameters and a minimum contig length of 2.5 kb. The depth profiles generated during the MetaBAT run, along with the BAM files, were subsequently used to perform binning with two additional tools: CONCOCT (v0.4.2, Alneberg et al., 2014) and MaxBin2 (v2.2.5., Wu et al., 2016). All bins from the three binners were refined and consolidated using MetaWRAP (v1.3-4bf5f8a, Uritskiy et al., 2018) in default mode.

To remove redundant genomes, the resulting MAGs were dereplicated at 99% Average Nucleotide Identity (ANI) using dRep (v2.3.2, Olm et al., 2017), yielding a final non-redundant set of 1,505 MAGs. For downstream analyses, only high-quality MAGs with >90% completeness and <10% contamination were selected, resulting in 461 high-quality MAGs used in this study.

### Dataset preparation and co-occurrence analyses

Before determining gene co-occurrence to determine potential metabolic interactions, individual genes were classified according to their metabolic pathways using the KEGG classification (https://www.genome.jp/kegg/pathway.html). After metabolism annotation, co-occurrence analysis was performed for the prokaryotic and eukaryotic annotated genes using the Maximal Information-based Nonparametric Exploration (MINE v2, Reshef et al., 2011) statistics as well as the maximal information coefficient (MIC) to characterize the degree of dependence between the different genes. MINE parameters were set to “equitability” for the best equitability of MICe, while the alpha, c, and cv values were set to default.

The resulting data was processed as follows: first, all negative associations (or edges) and those with a MIC below 0.5 were discarded. Negative associations were discarded due to their more complex and ambiguous interpretation. Unlike positive associations, which suggest shared niches or other ecological interactions, negative associations (typically reflecting a lack of co-occurrence) cannot be reliably interpreted as evidence of antagonism or competition without further ecological or experimental validation. After dataset curation, all KOs were clustered according to their KEGG metabolism, and the number of detected associations between different metabolisms was calculated using R (v4.3.1). As each metabolism included a different number of KOs, the number of edges was normalized by the number of KOs in both connected metabolisms in order to compare the density of associations between the different metabolisms. After normalization, we constructed a network to study the KO co-occurrence across the different metabolisms using Cytoscape (v3.10.1), and the network statistics were analyzed. In these networks, nodes represent KOs, while edges indicate significant associations between them.

To determine the edge density between prokaryotic vitamin biosynthesis and eukaryotic metabolisms, eukaryotic and prokaryotic KOs were differentiated before the co-occurrence analysis. After that, only the associations between eukaryotic and prokaryotic metabolisms with Cobalamin *de novo* biosynthesis (including aerobic and anaerobic pathways) and Thiamine *de novo* biosynthesis pathways (including all possible pathways) were studied. To enable comparison of association density across different metabolisms, the number of edges per metabolic pathway was normalized by the number of KOs within each pathway, following the same approach used in the previous analysis.

### MAG’s metabolic capabilities and co-occurence

Only MAGs with completeness above 90% and contamination below 10% were considered (totaling 461 MAGs). The potential ability to synthesize *de novo* cobalamin and thiamine was explored in these MAGs by comparing the completeness of the pathways for synthesizing both vitamins, looking at the presence of the different KOs according to the KEGG database. We classified a MAG as a complete cobalamin synthesizer if it contained more than 75% of the orthologs required for each step of cobalamin biosynthesis: corrinoid ring biosynthesis, nucleotide loop assembly, and lower ligand activation (via CobC and CobT enzymes). A MAG was categorized as a partial cobalamin synthesizer if it possessed more than 75% of the orthologs for corrinoid ring biosynthesis and nucleotide loop assembly but lacked the necessary genes for lower ligand activation. Finally, a MAG was designated as a lower ligand activator if it contained less than 75% of the orthologs for corrinoid ring biosynthesis and/or nucleotide loop assembly but retained the ability to activate the lower ligand.

Thiamine biosynthesis completeness was assessed based on the ability to synthesize its two main vitamers: 4-amino-5-hydroxymethyl-2-methylpyrimidine (HMP) and 4-methyl-5-(2-hydroxyethyl)thiazole (THZ). These compounds are produced through distinct pathways and subsequently joined by thiE to form thiamine. We classified pyrimidine auxotrophs as those MAGs that contain thiE and the thiG enzyme required for THZ synthesis but lack thiC, which is responsible for HMP synthesis. Thiazole auxotrophs were defined as MAGs possessing thiE and thiC but missing thiG, while dual auxotrophs were those containing thiE but lacking both thiC and thiG.

After determining the vitamin pathway completeness in each MAG, the co-occurrence analysis was performed with MINE using the same parameters as described earlier as well as the MAG’s abundances based on Reads Per Kilobase of Genome per Gigabase of Metagenome (RPKG). The network visualization and statistical analyses were performed in Cytoscape as described above. In the network, nodes represent individual MAGs, while edges indicate significant associations between them.

To determine the main core MAGs in the network, we used the approach described in Krabberød et al. (2022). For each MAG, we calculated 1) Degree: number of edges per node, 2) Betweenness centrality, which measures the frequency with which a MAG appears on the shortest paths between other MAGs in the network (meaning the most direct routes connecting two nodes in a network, using the fewest number of steps or edges); and 3) Closeness centrality, which indicates the proximity of a node to all other nodes in the network. We identify as “core” MAGs the ones that are within the first quartile for the three statistics. Among these, we defined the “main core MAGs” as those that ranked within the top 5% for each of the three centrality measures, indicating their central structural role in the network.

## RESULTS

### Environmental analysis

Between January 2009 and December 2020, the analysis of the time series (**Figure 1**) revealed seasonal variations in temperature, with maximum values recorded in August and September, peaking at 26.7 °C in August 2020, and minimum values observed between February and March, reaching a lowest temperature of 11.9 °C in March 2010. Chlorophyll *a* concentration also exhibited seasonal fluctuations, ranging from 0.1 to 2.9 µg l⁻¹, with peak values occurring in late winter and/or autumn. Inorganic nutrient concentrations showed distinct trends, with nitrate levels consistently peaking in winter, whereas ammonium and phosphate exhibited less predictable patterns.

**Figure 1.**
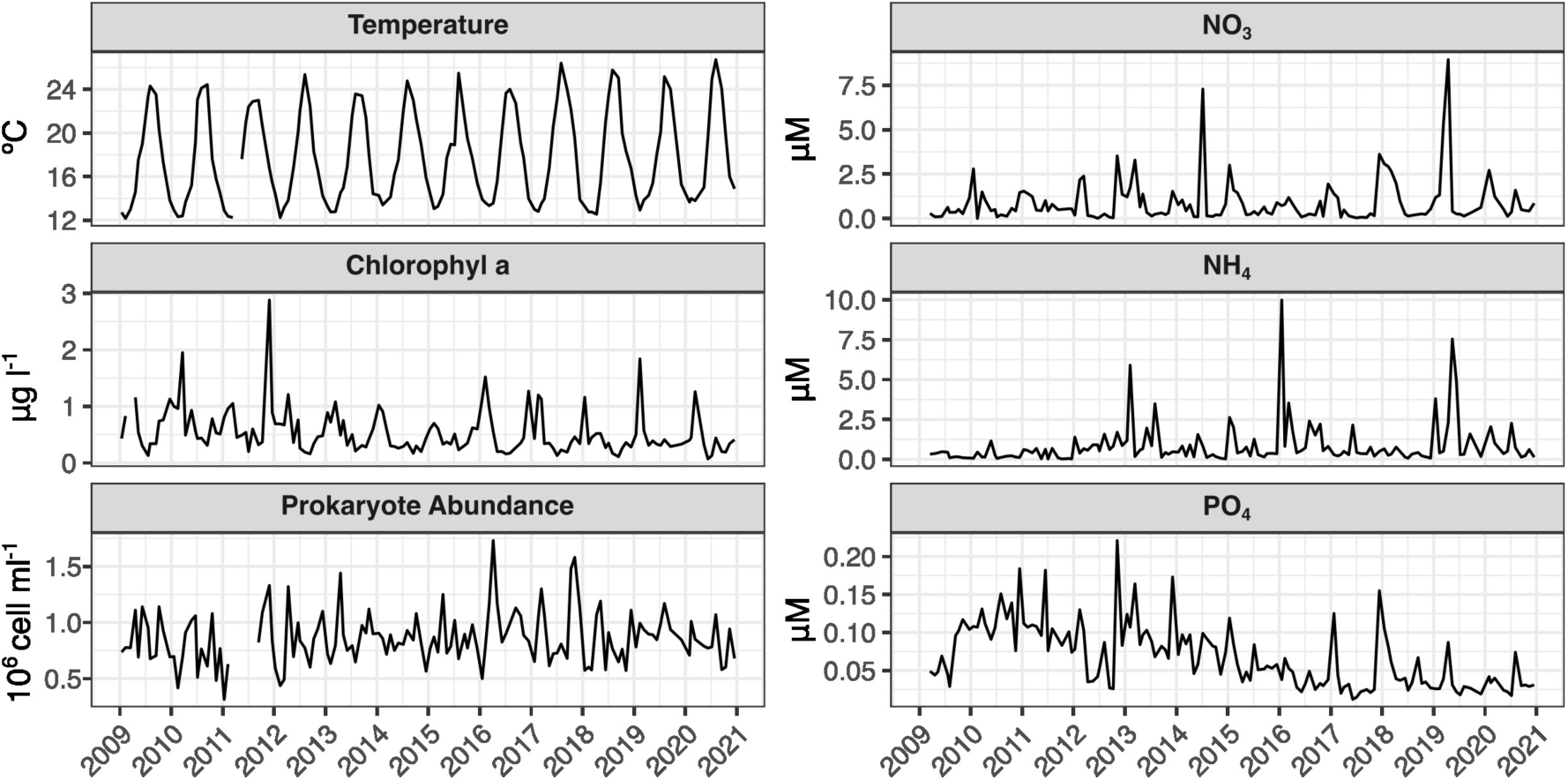
Temporal variation of environmental properties at the Blanes Bay Microbial Observatory (Catalonia) between 2009 and 2021. The figure shows monthly measurements of temperature, nitrate (NO₃⁻), ammonium (NH₄⁺), phosphate (PO₄³⁻), chlorophyll *a* concentration, and prokaryote abundance.

### Potential interactions among functions

Gene annotation identified a total of 8,370 prokaryotic KOs and 7,552 eukaryotic KOs. However, to filter the total number of KOs used for analysis, only the top 50% of the most abundant KOs from each group were used as input for the MINE analysis. The resulting network consisted of 2,225 nodes (representing KOs) and 2,377,290 edges (or associations). For further analysis, only positive associations with a maximal information coefficient (MIC) of 0.5 or higher were considered. Nodes were clustered according to the pathway and metabolic classifications assigned to each KO in the Kyoto Encyclopedia of Genes and Genomes (KEGG) database (ww.genome.jp/kegg/pathway.html), and their temporal associations were analyzed (Figure 2). The analysis revealed that pathways related to cofactor and vitamin metabolism exhibited the highest number of associations, with an average of 7.6 edges per node, followed closely by central carbohydrate metabolism (7.4 edges per node). Although many edges connected KOs within the same metabolic group, most vitamin metabolism-related KOs were associated with central carbohydrate metabolism, showing an average of 16.5 edges per node. Similarly, the metabolism of purine and amino acids (including lysine and histidine) played a central role in the network, with an average of 5.9 edges per node.

**Figure 2.**
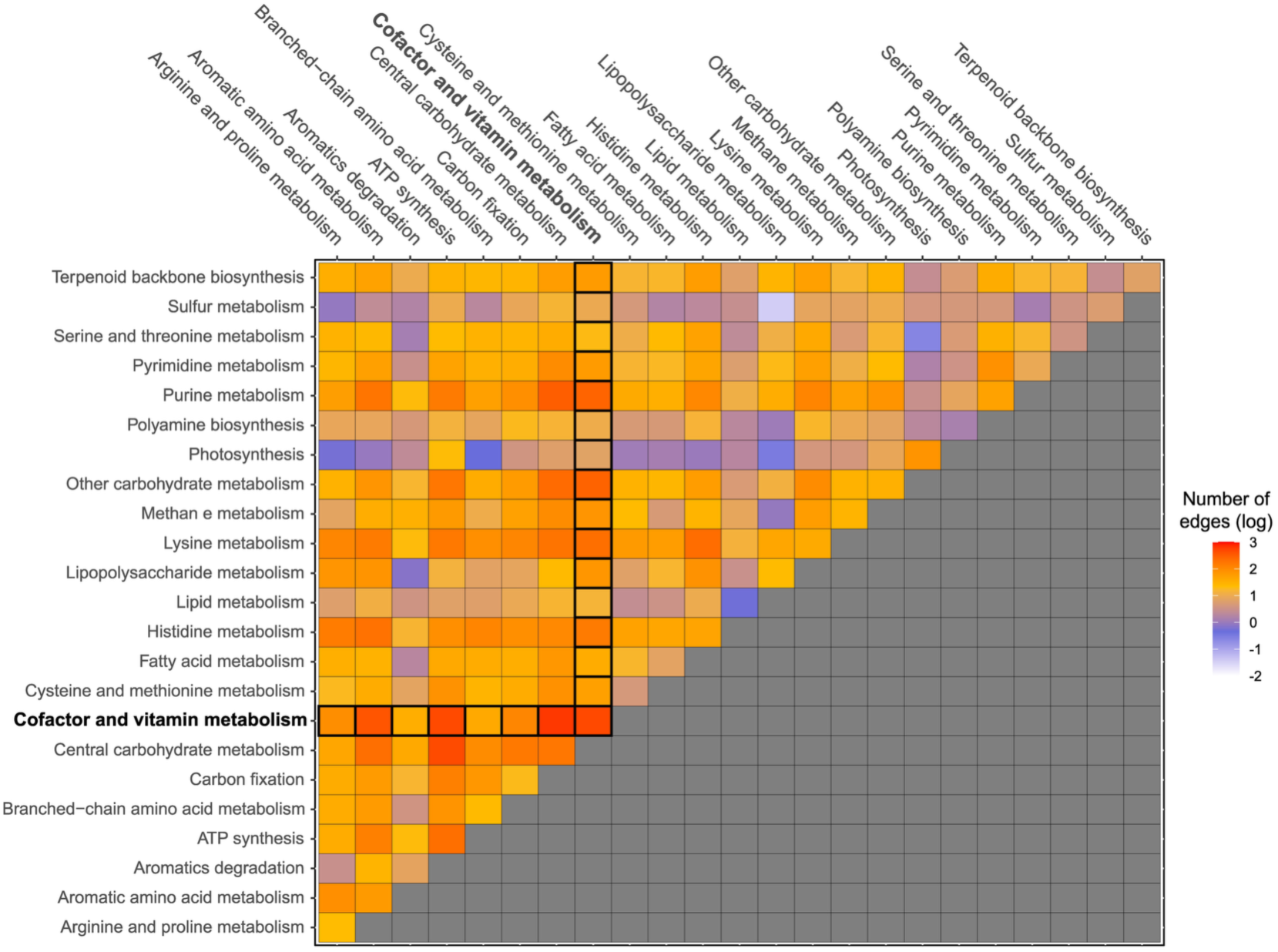
Heatmap comparing the number of edges (log scale) between KEGG metabolisms normalized by the number of KOs for each metabolism group. The Cofactor and Vitamin Metabolism group is highlighted.

When comparing the number of edges per node for prokaryotic vitamin biosynthesis-related KOs with other prokaryotic and eukaryotic metabolic pathways (Figure 3), we found that prokaryotic vitamin biosynthesis KOs exhibited the highest number of associations with eukaryotic metabolic pathways. Among these, eukaryotic central metabolic processes, particularly carbon fixation and carbohydrate metabolism, showed the strongest connectivity with prokaryotic vitamin biosynthesis KOs. Furthermore, when comparing the number of associations with the biosynthesis of individual vitamins (cobalamin, Thiamine, Biotin, and Riboflavin, Figure 3 and Supplementary Figure 2), we observed that cobalamin biosynthesis was more related to eukaryotic and prokaryotic metabolism (average 19.0 edges per node), followed by Thiamine (12.3 edges per node), while Biotin and Riboflavin showed 8.8 and 3.2 edges per node respectively. To further compare the association between prokaryotic and eukaryotic metabolisms with the biosynthesis of individual vitamins, we calculated the ratio of edge density between all the KOs involved in the prokaryotic biosynthesis of each vitamin with the KOs of the main central metabolisms for both prokaryotic and eukaryotic groups (Supplementary Figure 3). The results revealed stronger association metrics between prokaryotic vitamin synthesizers and eukaryotes than between prokaryotes themselves.

**Figure 3.**
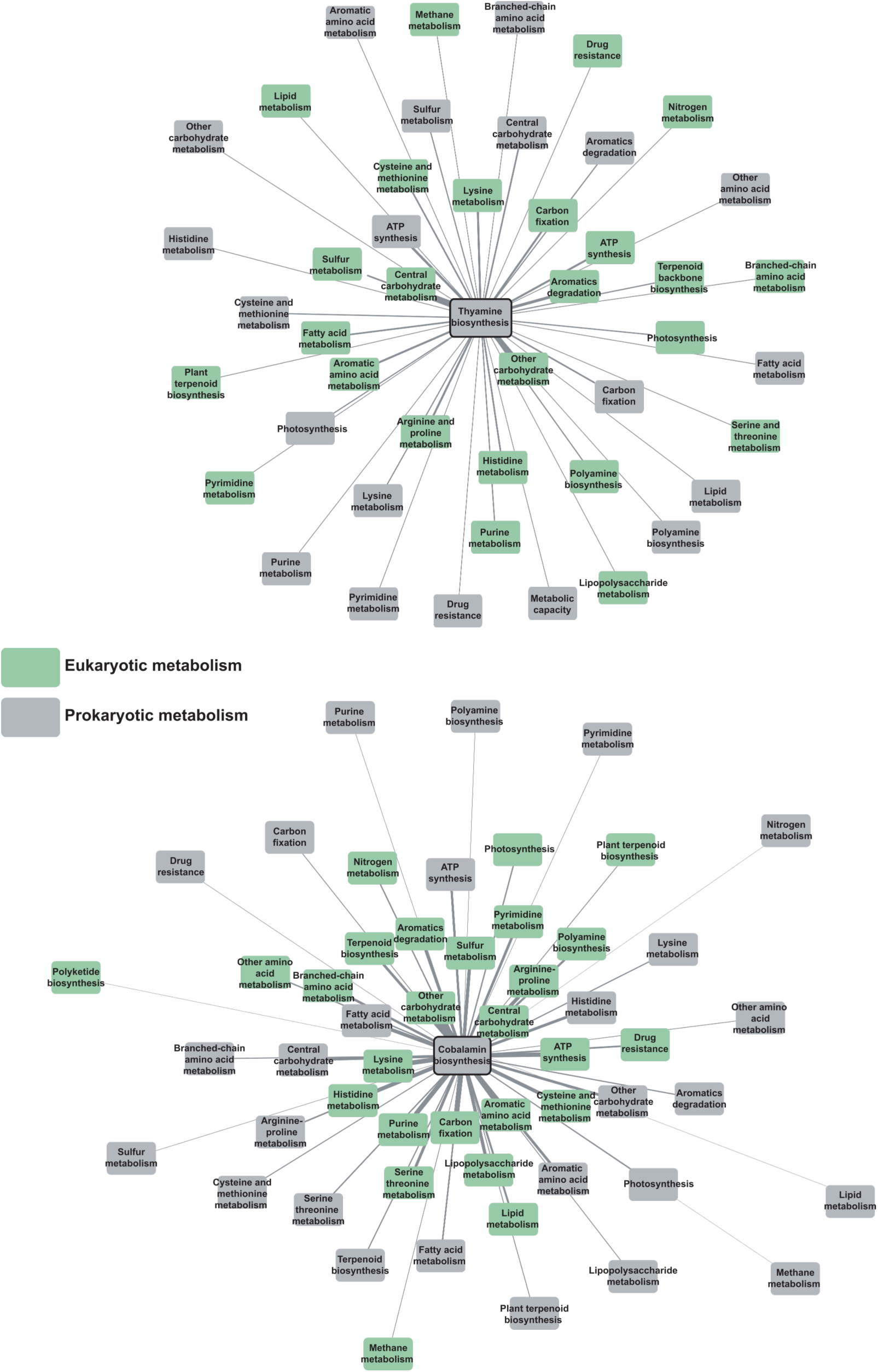
Co-occurrence networks generated with MINE, based on distinct KOs involved in the biosynthesis of thiamine (top) and cobalamin (bottom). All known biosynthetic pathways for each vitamin were considered for KO selection. Vitamin synthesis KOs are connected to various prokaryotic (grey) and eukaryotic (green) metabolic pathways. The proximity of the nodes to the central thiamine or cobalamin biosynthesis node reflects their number of connections, with closer nodes indicating higher edge density and thus greater connectivity within each metabolic context.

### MAG analysis and vitamin biosynthesis pathway completeness

Completeness of the *de novo* vitamin biosynthesis pathways was determined in recovered MAGs with genome completeness > 90%. A total of 461 MAGs were used, with an average genome completeness of 94.9%, an average contamination of 2%, and a GC content of 0.48. The average genome size per MAG was 3.12 Mb, and the number of predicted genes was 1,315,247 (average 2,853 genes per MAG). The selected MAGs represented, on average, around 10% of the total metagenome reads per sample (peaking at 36% in January 2011), while accounting for 23% of the abundance of all MAGs (Figure 4).

**Figure 4.**
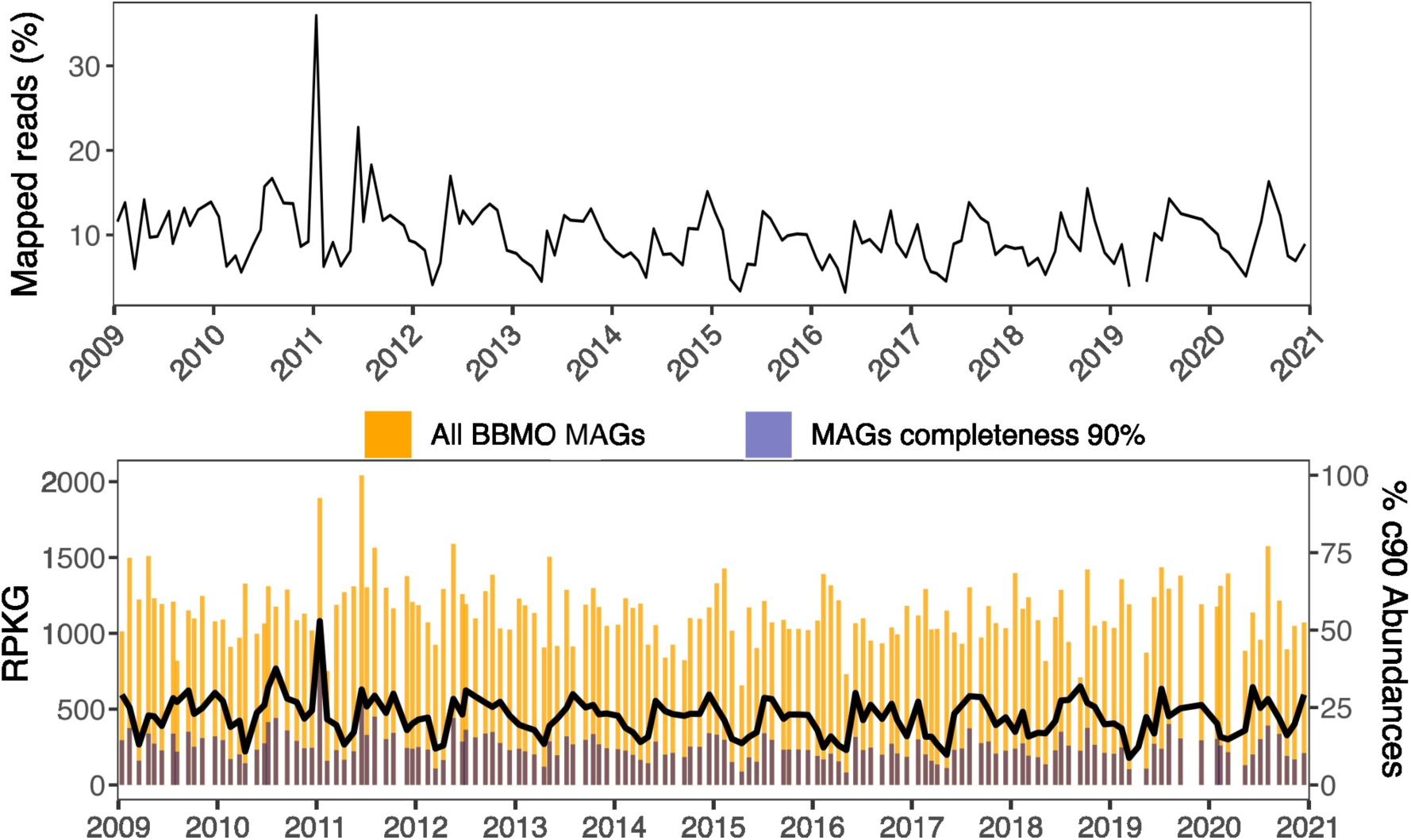
Metagenomic read recruitment and abundance of metagenome-assembled genomes (MAGs) at the Blanes Bay Microbial Observatory (BBMO) from 2009 to 2020. Top panel shows the percentage of metagenomic reads mapped to MAGs with ≥90% completeness. Bottom panel represents the total abundance of all MAGs (orange bars) and MAGs with ≥90% completeness (purple bars), expressed in RPKG (Reads Per Kilobase of genome per Gigabase of metagenome). The black line indicates the percentage of MAGs with ≥90% completeness relative to the total MAG abundance.

Overall, cobalamin de novo biosynthesis pathways exhibited the lowest completeness values, with only 11.5% of the MAGs (53 out of 461) containing a complete cobalamin biosynthesis pathway (Supplementary Figure 1). This includes the ability to synthesize the corrinoid ring through either aerobic or anaerobic pathways, assemble the nucleotide loop, and produce the complete lower ligand, requiring both CobC and CobT enzymes. In contrast, 2.8% of the MAGs were able to partial cobalamin biosynthesis (including aerobic or anaerobic corrinoid ring biosynthesis and nucleotide loop assembly but no ability for activation of the lower-ligand), and 11.1% of the MAGs were lower ligand activators (unable to synthesize the corrinoid ring and the nucleotide loop assembly, but able to activate the lower-ligand). Such a low abundance of cobalamin complete and partial synthesizers contrasts with the ability of almost the entire set of MAGs to synthesize Riboflavin (92.4%), and 40.8% of the MAGs showing at least one complete pathway for the *de novo* synthesis of Biotin.

As with cobalamin synthesizers, we observed the presence of both complete and partial thiamine synthesizers. Complete thiamine synthesizers, those with the *thiC*, *thiG* (or *thi4*), and *thiE* (or *thiN*) genes required to synthesize all thiamine vitamers (HMP-P, THZ-P, and their combination), constituted 32.5% of all MAGs. Partial synthesizers, capable of producing only the HMP or THZ precursors, represented 35.8% of the MAGs. Among these partial synthesizers, 36.2% lacked the *thiC* gene, making them pyrimidine B1 auxotrophs dependent on external pyrimidine. In contrast, 6.2% were thiazole auxotrophs, lacking *thiG* or *thi4*. Dual auxotrophs, lacking both pyrimidine synthase (*thiC*) and thiazole synthase (*thiG* or *thi4*) but retaining thiamine monophosphate synthase (*thiE* or *thiN*), accounted for 46.9% of MAGs.

The Rhodobacteraceae order was the most abundant group of cobalamin synthesizers, accounting for 36% of the total cobalamin prototrophic MAGs (Supplementary Figure 4). In contrast, the rest of the vitamin prototrophy exhibited greater diversity, with the Flavobacteriales order being the most abundant group. Specifically, Flavobacteriales represented 11.5% of Thiamine, 17.3% of Biotin, and 24% of Riboflavin prototrophs.

### Distribution of vitamin synthesizers

The co-occurrence analysis of the MAG dataset (including only the MAGs with a completeness > 90%) revealed two distinct subnetworks, each exhibiting clear seasonal differentiation (Figure 5). One subnetwork comprised 108 MAGs, which were almost exclusively present in the summer, while the other included 172 MAGs that appeared only during the winter. The taxonomical analysis of the MAGs revealed that the summer network was dominated by the Rhodobacterales and Puniceispirillales orders (25% of the MAGs each). The MAG with the highest abundance (RPKG) was G2.161 (Puniceispirillales order). On the other hand, we found a higher diversity in the winter network, with most MAGs belonging to the Rhodospirillales and Acidimicrobiales orders (11% each), although other relevant orders such as Synechococcales and Nitrososphaerales were also found (the one with the highest abundance (RPKG) was the archaeal MAG G3.122, of order Nitrososphaerales).

**Figure 5.**
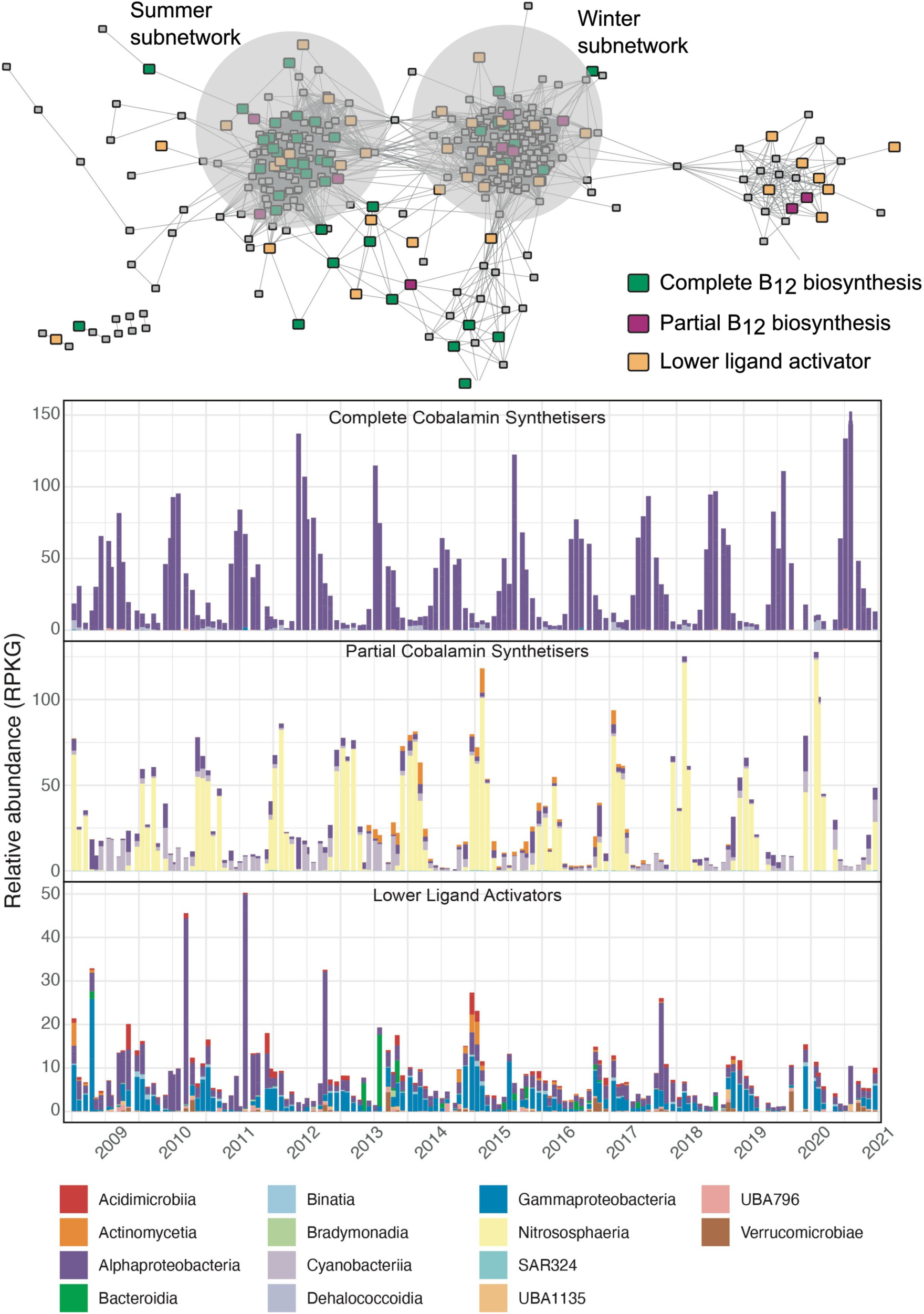
(Top) Association network of MAGs (>90% completeness, and <10% contamination), categorized by cobalamin biosynthesis completeness: complete (green), partial (purple), and lower ligand activators (orange). The summer and winter subnetworks are indicated by grey circles. (Bottom) Relative abundance of MAGs grouped at the class level based on their cobalamin biosynthesis classification: Complete Synthesizers, Partial Synthesizers, and Lower Ligan Activators.

The distribution of cobalamin and thiamine synthesizers (including both partial and complete synthesizers) was also analyzed to track vitamin-related interactions. Complete cobalamin synthesizers were present primarily during the summer, representing 20.5% of the summer subnetwork MAGs, and decreased to 6.7% in winter. This group was vastly dominated by Alphaproteobacteria, with Puniceispirillales and Rhodobacterales as the main families. On the other hand, partial cobalamin synthesizers and lower ligand activators were mainly found in the winter subnetwork (representing 3.0% and 8.5% of winter subnetwork MAGs, respectively). Nitrososphaeria dominated the partial synthesizer group, while the lower ligand activators were more diverse (Figure 5).

Regarding thiamine, complete synthesizers showed a higher prevalence in winter (40%) compared to summer (22.5%), while partial synthesizers were more equally distributed over seasons (28.4% in winter and 28.8% in summer, Supplementary Figure 5). However, pyrimidine auxotrophs and thiazole auxotrophs showed distinct seasonal distributions, with the former dominating in summer and the latter more prevalent in winter, largely due to the high abundance of the thiazole auxotrophic Nitrososphaeria group.

### Core MAGs in the co-occurrence network

To identify the key vitamin synthesizers connecting with other microbial groups, we analyzed network metrics, including degree (number of edges per node), betweenness centrality, and closeness centrality. We considered core nodes to be those within the top 25th percentile in all three statistics in both the summer and winter seasons (**Table 1**), as proposed by Krabberød et al. (2022). Additionally, we included more conservative percentiles (15th and 5th), with the most central nodes in each percentile determined by their betweenness centrality and closeness centrality. In winter, the most central node (within the 5th percentile) belonged to the Nitrososphaerales order (MAG.G3.122), while in summer, it was part of the Alphaproteobacteria class (Thalassobaculales order, MAG.G4.44). Interestingly, the two most central nodes in winter were both cobalamin synthesizers, either partial or complete. In summer, while complete cobalamin synthesizers comprised 20.5% of the MAGs, they accounted for 40% of the central nodes. In contrast, thiamine complete synthesizers were scarcely represented among the most central nodes in summer, with only a single MAG detected. However, their presence was markedly higher in winter, where they featured prominently on the central node list.

**Table 1.** Statistics of the most central nodes (MAGs) found in the Summer and Winter subnetworks calculated according to the Degree, Betweenness Centrality, and Closeness Centrality of each node (following the terminology in the Methods section and in Krabberød et al., 2022). Degree indicates the number of edges per node. Betweenness centrality, represents the most direct path connecting two nodes, using the fewest number of edges in between. Closeness centrality indicates the proximity of a node to all the other nodes in the network. For each MAG, the taxonomic affiliation and the ability to synthesize Cobalamin and Thiamine through different pathways are indicated.

| MAG | Degree | Betweenness Centrality | Closeness Centrality | Phylum | Class | Order | Core Percentile | Cobalamin synthesizer | Cob Lower Ligand | DMB Synthesis | Thiamine Complete | Thiamine cHET (thiG-thi4) | Thiamine HMP (thiC) | Thiamine Join (thiE) | Thiamine Savage |
| --- | --- | --- | --- | --- | --- | --- | --- | --- | --- | --- | --- | --- | --- | --- | --- |
| **Winter subnetwork** |  |  |  |  |  |  |  |  |  |  |  |  |  |  |  |
| bin.G3.122 | 145 | 0.0565 | 0.4707 | Thermoproteota | Nitrososphaeria | Nitrososphaerales | 5 | Yes |  | Yes |  |  | Yes |  |  |
| bin.G1.417 | 136 | 0.0890 | 0.4952 | Proteobacteria | Alphaproteobacteria | Bin95 | 15 | Yes | Yes |  |  |  |  | Yes |  |
| bin.G1.673 | 137 | 0.0336 | 0.4462 | Bacteroidota | Bacteroidia | Flavobacteriales | 15 |  |  |  | Yes |  |  |  |  |
| bin.G1.196 | 146 | 0.0098 | 0.4484 | Planctomycetota | Planctomycetes | Pirellulales | 15 |  |  |  |  | Yes | Yes |  |  |
| bin.G1.636 | 137 | 0.0087 | 0.4484 | Proteobacteria | Alphaproteobacteria | Rhodobacterales | 15 |  |  |  |  |  |  | Yes |  |
| bin.G1.687 | 132 | 0.0566 | 0.4634 | Bacteroidota | Bacteroidia | Flavobacteriales | 25 |  |  |  |  |  |  |  |  |
| bin.G1.90 | 133 | 0.0406 | 0.4769 | Myxococcota | Myxococcales | UBA4151 | 25 |  |  |  | Yes |  |  |  |  |
| bin.G3.280 | 129 | 0.0193 | 0.4529 | Bacteroidota | Bacteroidia | Cytophagales | 25 |  |  |  |  |  |  |  |  |
| bin.G1.556 | 129 | 0.0082 | 0.4392 | Proteobacteria | Alphaproteobacteria | UBA6615 | 25 | Yes | Yes |  |  |  |  | Yes |  |
| bin.G1.604 | 130 | 0.0061 | 0.4257 | Myxococcota | Myxococcales | UBA796 | 25 |  |  |  | Yes |  |  |  |  |
| bin.G1.343 | 140 | 0.0047 | 0.4349 | Verrucomicrobiota | Verrucomicrobiae | Opitutales | 25 |  |  |  | Yes |  |  |  |  |
| bin.G1.115 | 139 | 0.0044 | 0.4529 | Chloroflexota | Dehalococcoidia | SAR202 | 25 |  |  |  |  | Yes |  | Yes |  |
| bin.G1.220 | 135 | 0.0037 | 0.4397 | Actinobacteriota | Acidimicrobiia | Acidimicrobiales | 25 |  |  |  |  |  |  |  |  |
| bin.G1.841 | 146 | 0.0036 | 0.4392 | Latescibacterota | Latescibacteriia | UBA2968 | 25 |  |  |  |  |  |  |  |  |
| bin.G1.259 | 146 | 0.0035 | 0.4457 | Actinobacteriota | Acidimicrobiia | Acidimicrobiales | 25 |  |  | Yes |  |  |  |  |  |
| bin.G1.709 | 143 | 0.0034 | 0.4501 | Bacteroidota | Bacteroidia | Flavobacteriales | 25 |  |  |  |  | Yes |  | Yes |  |
| **Summer subnetwork** |  |  |  |  |  |  |  |  |  |  |  |  |  |  |  |
| bin.G4.44 | 78 | 0.0522 | 0.4038 | Proteobacteria | Alphaproteobacteria | Thalassobaculales | 5 | Yes | Yes |  |  |  |  |  |  |
| bin.G2.106 | 80 | 0.0398 | 0.4011 | Bacteroidota | Bacteroidia | Flavobacteriales | 5 |  |  |  |  |  |  |  |  |
| bin.G1.761 | 77 | 0.0973 | 0.4429 | Bacteroidota | Bacteroidia | Chitinophagales | 15 |  |  |  |  |  |  |  |  |
| bin.G3.433 | 73 | 0.0275 | 0.3967 | Proteobacteria | Alphaproteobacteria | Rhodobacterales | 15 | Yes | Yes |  |  |  |  | Yes |  |
| bin.G4.48 | 77 | 0.0093 | 0.3658 | Proteobacteria | Alphaproteobacteria | Rhodobacterales | 15 | Yes | Yes |  |  |  |  |  |  |
| bin.G2.161 | 74 | 0.0084 | 0.3665 | Proteobacteria | Alphaproteobacteria | Puniceispirillales | 15 |  |  |  |  |  |  | Yes | Yes |
| bin.G4.170 | 73 | 0.0079 | 0.3614 | Verrucomicrobiota | Verrucomicrobiae | Opitutales | 25 |  |  |  |  |  |  |  |  |
| bin.G4.400 | 71 | 0.0065 | 0.3632 | Proteobacteria | Alphaproteobacteria | UBA8317 | 25 |  |  |  |  |  |  | Yes | Yes |
| bin.G1.780 | 76 | 0.0062 | 0.3665 | Proteobacteria | Alphaproteobacteria | Puniceispirillales | 25 | Yes | Yes |  |  | Yes |  | Yes |  |
| bin.G1.286 | 79 | 0.0059 | 0.3643 | Proteobacteria | Alphaproteobacteria | Parvibaculales | 25 |  |  |  | Yes |  |  |  |  |

## DISCUSSION

The seasonal taxonomic co-occurrence among marine microbes has been extensively explored in several time series study sites like the San Pedro Ocean Time series station (SPOT, Steel et al., 2011; Chow et al., 2014; Cram et al., 2015) or the BBMO (Giner et al., 2019; Krabberød et al., 2022; Auladell et al., 2022; Deutschmann et al., 2023), elucidating the main associations among taxa (including prokaryotes and eukaryotes). While these studies have contributed significantly to our understanding of microbial interactions, relying solely on taxonomic co-occurrence may limit insights into the functional bases of specific microbial interactions. By contrast, our approach leverages gene and MAG co-occurrence, providing a more nuanced and functionally relevant perspective on metabolic microbial interactions. Recent research employing ecosystem modeling and genome-resolved metagenomics has underscored the critical role of microbial auxotrophies, particularly amino acid and vitamin B auxotrophies, in shaping microbial dynamics in the global ocean (Giordano et al., 2024). Our observations support the central role of vitamin-related metabolism in marine microbial interactions, as evidenced by the higher number of associations involving vitamin metabolism-related genes compared to other metabolic pathways; in agreement with previous studies (Croft et al., 2005; Kazamia et al., 2012; Sultana et al., 2023). Furthermore, our findings emphasize that the seasonal variation in the genomic capacity for vitamin biosynthesis (reflected in the distinct temporal distribution of complete and partial vitamin synthesizers) has the potential to influence the temporal interaction dynamics of prokaryotic and eukaryotic assemblages.

Vitamins are typically present at extremely low concentrations in marine systems, often in the order of picomolar (pM). Vitamin concentrations in the Mediterranean Sea remain at low pM level (Bonnet et al., 2013; Suffridge et al., 2018), suggesting a potential limitation for microbial activity. Earlier studies suggested that trace-level vitamin concentrations are sufficient to sustain microbial growth, since vitamins, as cofactors, are needed in low amounts and can be recycled (Droop et al., 2007; Paerl et al., 2018). Yet recent growth studies have shown that vitamins are often present at growth-limiting concentrations for auxotrophic microorganisms in the global oceans (Gregor et al., 2023). Therefore, de novo vitamin B synthesis is necessary to meet the vitamin requirements of many eukaryotes and prokaryotes in oligotrophic regions. Yet, despite the widespread presence of several vitamin biosynthesis genes in the metagenomes, microbes with complete vitamin biosynthesis pathways are scarce, most likely due to the high complexity of vitamin *de novo* synthesis (particularly for cobalamin, for which more than 30 genes are required for its complete synthesis). Our results align with previous observations (Sañudo-Wilhelmy et al., 2014) showing that approximately 41% of the microbial population possess a complete (or nearly complete, >75%) pathway for Biotin biosynthesis, while 32.5% of the species exhibited the capability to synthesize Thiamine *de novo* (Supplementary Figure 1). In contrast, the overall low abundance of MAGs with a complete pathway for the *de novo* synthesis of cobalamin (a vitamin required by most prokaryotic and half of the eukaryotic marine microorganisms; Sañudo-Wilhelmy et al., 2014; Zoccarato et al., 2022) highlights the imbalance between B_12_ producers and auxotrophs. This scarcity likely drives the higher number of associations (edges) per node observed between central metabolic pathways and B_12_ synthesis genes compared to those for B_1_ or B_7_ biosynthesis.

It is important to acknowledge that our reliance on complete MAGs to resolve the complex steps of vitamin biosynthesis limits our analysis to only a subset of the microbial community. Nonetheless, the strong seasonal trends of MAGs observed here closely align with those reported in a recent study on vitamin biosynthesis genes in the Mediterranean Sea (Beauvais et al., 2023). Moreover, the proportion of vitamin synthesizers identified in our dataset is consistent with previous observations (Sañudo-Wilhelmy et al., 2012), suggesting that our findings provide a representative picture of the vitamin-related interactions occurring at the community level.

The recently described phenomenon of cobalamin lower ligand cross-feeding likely contributes to the high number of B_12_-related correlations observed in our network analyses (Wienhausen et al., 2024). Cobalamin biosynthesis requires the assembly of three key components: the corrin ring, the nucleotide loop, and the activated form of the lower ligand (alpha-ribazole). The lower ligand, typically 5,6-dimethylbenzimidazole (DMB) and catalyzed by the bluB enzyme, is essential for cobalamin activation and is incorporated through the action of CobC and CobT enzymes, which convert it into its activated form, α-ribazole. Wienhausen et al. (2024) demonstrated that the lower ligand can be exchanged between microorganisms through cross-feeding, compensating for the limited number of microbes with a complete cobalamin biosynthesis pathway. The strong associations between partial synthesizers and lower-ligand activators, which exhibit the highest edge density between both groups (see Supplementary Table 1), further support the idea that ligand cross-feeding likely represents an important microbial interaction throughout the year. However, the pronounced seasonal variation between complete and partial synthesizers suggests that cross-feeding interactions are more prevalent in winter. Notably, the number of associations between partial synthesizers and lower-ligand activators was significantly higher in winter (44 edges) compared to summer (2 edges), strongly indicating a seasonal shift in cobalamin biosynthesis strategies.

However, the bluB enzyme, responsible for the production of DMB (Gray & Escalante-Semerena, 2007; Wienhausen et al., 2024), was found in only a small fraction of MAGs: 7% of MAGs containing CobC and CobT enzymes and just 12 MAGs with complete cobalamin biosynthesis pathways. Remarkably, nearly half of the MAGs encoding bluB are incapable of synthesizing any cobalamin-related pathways (complete, partial, or lower-ligand activation), suggesting that these microbes might primarily contribute to DMB production for cross-feeding. Seasonal trends further highlight the role of <u>bluB</u>, with 46% of MAGs containing bluB exclusive to the winter subnetwork compared to only 13% in summer. These findings support our previous findings that cross-feeding interactions become essential for achieving complete cobalamin synthesis during winter when fully independent synthesizers are less prevalent.

An example of a potential ligand cross-feeding interaction in winter is observed between the *Nitrosopumilus* (MAG bin.G3.122), the main central core of the winter subnetwork (Table 1), and different Lower-Ligand Activator MAGs. Previous studies have suggested that members of *Nitrosopumilus* can serve as an important source of B_12_ (Doxey et al., 2015; Qin et al., 2017; Santoro et al., 2015). However, our study reveals that *Nitrosopumilus*, despite being the dominant partial B_12_ producer in winter (accounting for up to 10% of total metagenome reads in winter), lacks the ability to activate the lower ligand. Interestingly, it retains the capacity to synthesize DMB through the presence of the *bluB* enzyme. This observation is further supported by available metagenomic data confirming the absence of CobC and CobT in *Nitrosopumilus* (Thaumarchaeota) genomes (Santoro et al., 2015), indicating the inability of this group to activate the lower ligand independently. Consequently, *Nitrosopumilus* could provide DMB to the environment but requires interactions with Lower-Ligand Activators to complete cobalamin biosynthesis. Our findings suggest that *Verrucomicrobia* (order Opitutales, bin.G4.517), one of the main groups in winter (Krabberød et al., 2022), and *Puniceispirillales* (bin.G1.296) are likely key contributors in this cross-feeding process, as they show a very close association with the *Nitrosopumilus* MAG (MIC 0.64 and 0.82). However, the relative abundance of Lower-Ligand Activators exhibited high interannual variability, likely driven by their greater diversity compared to partial synthesizers. This variability resulted in shifts in the dominant Lower-Ligand Activator groups over the years, with different members of *Alphaproteobacteria*, *Gammaproteobacteria*, *Bacteroidetes*, and even *Actinomycetia* taking on the role. This suggests that functional redundancy within the microbial community allows different taxa to compensate for shifts in group dominance (Ramond et al., 2025), maintaining the essential cross-feeding interactions needed for cobalamin biosynthesis in winter.

In contrast to cobalamin biosynthesis-related associations, *de novo* thiamine biosynthesis exhibited a significantly lower number of associations, likely reflecting the reduced need for exchanges to meet thiamine requirements due to the higher number of microorganisms capable of synthesizing thiamine (with 2.6 times more complete thiamine synthesizers compared to cobalamin). Despite the overall lower number of associations, thiamine synthesizers seem relevant in the winter network, with 4 of the 15 most central nodes in the winter subnetwork identified as Complete Thiamine Synthesizers. Such relevance in winter could be related to thiamine’s role as a cofactor in the TCA cycle, essential for metabolizing organic compounds such as carbohydrates and proteins (Rapala-Kozik, 2011). The BBMO site is characterized by a seasonal chlorophyll pattern, peaking in late winter (Gasol et al., 2016) due to blooms of diatoms and autotrophic picoplankton (Nunes et al., 2018; Vallina et al., 2023). This abundance of autotrophs likely increases the availability of organic molecules, stimulating microbial activity and respiration (García-Martín et al., 2015) and thus raising the demand for thiamine, which could explain the observed peak in thiamine-related interactions during winter.

For thiamine, maximum edge density (indicating a preferential association) was observed between complete thiamine synthesizers and auxotrophs (Supplementary Figure 1), highlighting the potential dependence of certain taxa on vitamin producers. Interestingly, thiazole auxotrophs also showed the highest edge density with pyrimidine auxotrophs, suggesting that, similar to cobalamin, cross-feeding interactions involving thiamine precursors may become particularly relevant when complete synthesizers are scarce. The overall higher prevalence of pyrimidine auxotrophy aligns with observations from the Baltic Sea (Paerl et al., 2018), while its increased occurrence in summer further suggests elevated environmental levels of the HMP during this period, consistent with observations from other marine environments (Bittner et al., 2024). Conversely, thiazole auxotrophs, mostly *Nitrosopumilus,* may utilize HET released by some cells of the highly abundant complete thiamine synthesizers during winter, reducing its dependency on other synthesizers to meet their B1 requirements.

Taken together, our findings provide new insights into the seasonal dynamics of vitamin biosynthesis and its role in structuring microbial interactions in marine ecosystems, particularly in relation to cobalamin and thiamine. By leveraging gene co-occurrence rather than solely relying on taxonomic data, our results suggest that complete and partial vitamin synthesis are among the main microbial interactions throughout the year. Based on experimental approaches of previous studies, our results highlight the critical importance of cross-feeding interactions, particularly for cobalamin, where lower ligand activation and seasonal succession in the microbial assemblage composition play a key role in maintaining vitamin availability. The distinct seasonal strategies on the biosynthesis of vitamins, with partial and complete synthesizers showing a different temporal distribution, highlight the dynamic nature of the vitamin-related microbial interactions.

Yet, the consistent seasonal pattern of vitamin biosynthesis genes contrasts with the interannual taxonomic shifts among vitamin producers, suggesting that functional redundancy (particularly in lower ligand activation) helps maintain key metabolic functions across the 12-year time series. Altogether, our results highlight vitamins as one of the main drivers of ecological interactions between eukaryotic and prokaryotic microorganisms in the Mediterranean Sea.

## Acknowledgement

We thank all members of the Blanes Bay Microbial Observatory (BBMO) team for their collaboration in the sample collection. We are also grateful to the multiple projects that fund the BBMO initiative (https://bbmo.icm.csic.es). The analyses of the project have been performed at the Marbits bioinformatics core at ICM-CSIC (https://marbits.icm.csic.es) and at the Supercomputing Center of Galicia (CESGA). This work has been partially funded by the MAORI project (PID2022-136281NB-I00). NAG was supported by the Beatriu de Pinós program (2020-BP-00179). This work acknowledges the Severo Ochoa Centre of Excellence accreditation (CEX2019-000928-S) funded by AEI 10.13039/501100011033.

## Notes

### Competing Interest Statement

The authors have declared no competing interest.

## References

Alneberg, J., Bjarnason, B. S., De Bruijn, I., Schirmer, M., Quick, J., Ijaz, U. Z., Lahti, L., Loman, N. J., Andersson, A. F., & Quince, C. (2014). Binning metagenomic contigs by coverage and composition. Nature Methods, 11(11), 1144–1146. 10.1038/nmeth.3103

Anders, S., Pyl, P. T., & Huber, W. (2015). HTSeq—A Python framework to work with high-throughput sequencing data. Bioinformatics, 31(2), 166–169. 10.1093/bioinformatics/btu638

Arandia-Gorostidi, N., Krabberød, A. K., Logares, R., Deutschmann, I. M., Scharek, R., Morán, X. A. G., González, F., & Alonso-Sáez, L. (2022). Novel Interactions Between Phytoplankton and Bacteria Shape Microbial Seasonal Dynamics in Coastal Ocean Waters. Frontiers in Marine Science, 9, 901201. 10.3389/fmars.2022.901201

Auladell, A., Barberán, A., Logares, R., et al. (2022). Seasonal niche differentiation among closely related marine bacteria. ISME Journal, 16, 178–189. 10.1038/s41396-021-01053-2

Beauvais, M., Schatt, P., Montiel, L., Logares, R., Galand, P. E., & Bouget, F. (2023). Functional redundancy of seasonal vitamin B_12_ biosynthesis pathways in coastal marine microbial communities. Environmental Microbiology, 25(12), 3753–3770. 10.1111/1462-2920.16545

Benoit, G., Peterlongo, P., Mariadassou, M., Drezen, E., Schbath, S., Lavenier, D., & Lemaitre, C. (2016). Multiple comparative metagenomics using multiset *k*-mer counting. PeerJ Computer Science, 2, e94. 10.7717/peerj-cs.94

Berg, J. M., Tymoczko, J. L., & Stryer, L. (2007). *Biochemistry* (Sixth ed). W. H. Freeman and company.

Bittner, M. J., Bannon, C. C., Rowland, E., Sundh, J., Bertrand, E. M., Andersson, A. F., Paerl, R. W., & Riemann, L. (2024). New chemical and microbial perspectives on vitamin B1 and vitamer dynamics of a coastal system. ISME Communications, 4(1), ycad016. 10.1093/ismeco/ycad016

Bonnet, S., Tovar-Sánchez, A., Panzeca, C., Duarte, C. M., Ortega-Retuerta, E., & Sañudo-Wilhelmy, S. A. (2013). Geographical gradients of dissolved Vitamin B12 in the Mediterranean Sea. Frontiers in Microbiology, 4. 10.3389/fmicb.2013.00126

Buchfink, B., Xie, C., & Huson, D. H. (2015). Fast and sensitive protein alignment using DIAMOND. Nature Methods, 12(1), 59–60. 10.1038/nmeth.3176

Chow, C.-E. T., Kim, D. Y., Sachdeva, R., Caron, D. A., & Fuhrman, J. A. (2014). Top-down controls on bacterial community structure: Microbial network analysis of bacteria, T4-like viruses and protists. The ISME Journal, 8(4), 816–829. 10.1038/ismej.2013.199

Combs, G. F., & McClung, J. P. (2017). The vitamins: Fundamental aspects in nutrition and health (Fifth edition). Elsevier/AP.

Cram, J. A., Xia, L. C., Needham, D. M., Sachdeva, R., Sun, F., & Fuhrman, J. A. (2015). Cross-depth analysis of marine bacterial networks suggests downward propagation of temporal changes. The ISME Journal, 9(12), 2573–2586. 10.1038/ismej.2015.76

Crocker, K., Lee, K. K., Chakraverti-Wuerthwein, M., Li, Z., Tikhonov, M., Mani, M., Gowda, K., & Kuehn, S. (2024). Environmentally dependent interactions shape patterns in gene content across natural microbiomes. Nature Microbiology, 9(8), 2022–2037. 10.1038/s41564-024-01752-4

Croft, M. T., Lawrence, A. D., Raux-Deery, E., Warren, M. J., & Smith, A. G. (2005). Algae acquire vitamin B12 through a symbiotic relationship with bacteria. Nature, 438(7064), 90–93.

Croft, M. T., Warren, M. J., & Smith, A. G. (2006). Algae Need Their Vitamins. Eukaryotic Cell, 5(8), 1175–1183. 10.1128/EC.00097-06

Danecek, P., Bonfield, J. K., Liddle, J., Marshall, J., Ohan, V., Pollard, M. O., Whitwham, A., Keane, T., McCarthy, S. A., Davies, R. M., & Li, H. (2021). Twelve years of SAMtools and BCFtools. GigaScience, 10(2), giab008. 10.1093/gigascience/giab008

Datta, M. S., Sliwerska, E., Gore, J., Polz, M. F., & Cordero, O. X. (2016). Microbial interactions lead to rapid micro-scale successions on model marine particles. Nature Communications, 7(1), 11965. 10.1038/ncomms11965

Deutschmann, I. M., Delage, E., Giner, C. R., Sebastián, M., Poulain, J., Arístegui, J., Duarte, C. M., Acinas, S. G., Massana, R., Gasol, J. M., Eveillard, D., Chaffron, S., & Logares, R. (2024). Disentangling microbial networks across pelagic zones in the tropical and subtropical global ocean. Nature Communications, 15(1). 10.1038/s41467-023-44550-y

Deutschmann, I. M., Krabberød, A. K., Latorre, F., Delage, E., Marrasé, C., Balagué, V., Gasol, J. M., Massana, R., Eveillard, D., Chaffron, S., & Logares, R. (2023). Disentangling temporal associations in marine microbial networks. Microbiome, 11(1), 83. 10.1186/s40168-023-01523-z

Dowling, D. P., Croft, A. K., & Drennan, C. L. (2012). Radical Use of Rossmann and TIM Barrel Architectures for Controlling Coenzyme B_12_ Chemistry. Annual Review of Biophysics, 41(1), 403–427. 10.1146/annurev-biophys-050511-102225

Doxey, A. C., Kurtz, D. A., Lynch, M. D. J., Sauder, L. A., & Neufeld, J. D. (2015). Aquatic metagenomes implicate Thaumarchaeota in global cobalamin production. ISME Journal, 9(2), 461–471. 10.1038/ismej.2014.142

Droop, M. R. (2007). Vitamins, phytoplankton and bacteria: Symbiosis or scavenging? Journal of Plankton Research, 29(2), 107–113. 10.1093/plankt/fbm009

Durham, B. P. (2021). Deciphering metabolic currencies that support marine microbial networks. mSystems, 6, e00763–21. 10.1128/mSystems.00763-21

Durham, B. P., Sharma, S., Luo, H., Smith, C. B., Amin, S. A., Bender, S. J., Dearth, S. P., Van Mooy, B. A. S., Campagna, S. R., Kujawinski, E. B., Armbrust, E. V., & Moran, M. A. (2015). Cryptic carbon and sulfur cycling between surface ocean plankton. Proceedings of the National Academy of Sciences, 112(2), 453–457. 10.1073/pnas.1413137112

Fondi, M., & Di Patti, F. (2019). A synthetic ecosystem for the multi-level modelling of heterotroph-phototroph metabolic interactions. Ecological Modelling, 399, 13–22. 10.1016/j.ecolmodel.2019.02.012

Fondi, M., Karkman, A., Tamminen, M. V., Bosi, E., Virta, M., Fani, R., Alm, E., & McInerney, J. O. (2016). “Every gene is everywhere but the environment selects”: Global geolocalization of gene sharing in environmental samples through network analysis. Genome Biology and Evolution, 8(5), 1388–1400. 10.1093/gbe/evw077

García-Martín, E. E., Daniels, C. J., Davidson, K., Lozano, J., Mayers, K. M. J., McNeill, S., Mitchell, E., Poulton, A. J., Purdie, D. A., Tarran, G. A., Whyte, C., & Robinson, C. (2019). Plankton community respiration and bacterial metabolism in a North Atlantic Shelf Sea during spring bloom development (April 2015). Progress in Oceanography, 177, 101873. 10.1016/j.pocean.2017.11.002

Gasol, J. M., Cardelús, C., Morán, X. A. G., Balagué, V., Forn, I., Marrasé, C., Massana, R., Pedrós-Alió, C., Sala, M. M., Simó, R., Vaqué, D., & Estrada, M. (2016). Seasonal patterns in phytoplankton photosynthetic parameters and primary production at a coastal NW Mediterranean site. Scientia Marina, 80(S1), 63–77. 10.3989/scimar.04480.06E

Gemayel, K., Lomsadze, A., & Borodovsky, M. (2022). MetaGeneMark-2: Improved Gene Prediction in Metagenomes. Bioinformatics. 10.1101/2022.07.25.500264

Gerald F., Combs, J., McClung, J. P., & McClung, J. P. (2017). *Vitamins: Fundamental Aspects in Nutrition and Health* (5th ed). Elsevier Science & Technology.

Giner, C. R., Balagué, V., Krabberød, A. K., Ferrera, I., Reñé, A., Garcés, E., Gasol, J. M., Logares, R., & Massana, R. (2019). Quantifying long-term recurrence in planktonic microbial eukaryotes. Molecular Ecology, 28(5), 923–935. 10.1111/mec.14929

Giordano, N., Gaudin, M., Trottier, C., et al. (2024). Genome-scale community modelling reveals conserved metabolic cross-feedings in epipelagic bacterioplankton communities. Nature Communications, 15, 2721. 10.1038/s41467-024-46374-w

Giovannoni, S. J. (2012). Vitamins in the sea. Proceedings of the National Academy of Sciences, 109(35), 13888–13889. 10.1073/pnas.1211722109

Gralka, M., Szabo, R., Stocker, R., & Cordero, O. X. (2020). Trophic Interactions and the Drivers of Microbial Community Assembly. Current Biology, 30(19), R1176–R1188. 10.1016/j.cub.2020.08.007

Graßhoff, K., Graßhoff, K., Kremling, K., & Ehrhardt, M. (Eds.). (2009). Methods of Seawater Analysis (3. vollst. überarb. u. erw. Auflage). Wiley-VCH.

Gray, M. J., & Escalante-Semerena, J. C. (2007). Single-enzyme conversion of FMNH2 to 5,6-dimethylbenzimidazole, the lower ligand of B12. *Proceedings of the National Academy of Sciences*, U.S.A., 104(8), 2921–2926. 10.1073/pnas.0609270104

Gregor, R., Vercelli, G. T., Szabo, R. E., Gralka, M., Reynolds, R. C., Qu, E. B., Levine, N. M., & Cordero, O. X. (2023). Vitamin auxotrophies shape microbial community assembly in the ocean. Microbiology. 10.1101/2023.10.16.562604

Hibbing, M. E., Fuqua, C., Parsek, M. R., & Peterson, S. B. (2010). Bacterial competition: Surviving and thriving in the microbial jungle. Nature Reviews Microbiology, 8(1), 15–25. 10.1038/nrmicro2259

Hyatt, D., Chen, G.-L., LoCascio, P. F., Land, M. L., Larimer, F. W., & Hauser, L. J. (2010). Prodigal: Prokaryotic gene recognition and translation initiation site identification. BMC Bioinformatics, 11(1), 119. 10.1186/1471-2105-11-119

Kanehisa, M. (2000). KEGG: Kyoto Encyclopedia of Genes and Genomes. Nucleic Acids Research, 28(1), 27–30. 10.1093/nar/28.1.27

Kang, D. D., Li, F., Kirton, E., Thomas, A., Egan, R., An, H., & Wang, Z. (2019). MetaBAT 2: An adaptive binning algorithm for robust and efficient genome reconstruction from metagenome assemblies. PeerJ, 7, e7359. 10.7717/peerj.7359

Kazamia, E., Czesnick, H., Nguyen, T. T. V., Croft, M. T., Sherwood, E., Sasso, S., et al. (2012). Mutualistic interactions between vitamin B12-dependent algae and heterotrophic bacteria exhibit regulation. Environmental Microbiology, 14(6), 1466– 1476.

Krabberød, A. K., Deutschmann, I. M., Bjorbækmo, M. F. M., et al. (2022). Long-term patterns of an interconnected core marine microbiota. Environmental Microbiome, 17, 22. 10.1186/s40793-022-00417-1

Kuppa Baskaran, D. K., Umale, S., Zhou, Z., Raman, K., & Anantharaman, K. (2023). Metagenome-based metabolic modelling predicts unique microbial interactions in deep-sea hydrothermal plume microbiomes. ISME Communications, 3(1), 42. 10.1038/s43705-023-00242-8

Li, D., Liu, C. M., Luo, R., Sadakane, K., & Lam, T. W. (2015). MEGAHIT: An ultra-fast single-node solution for large and complex metagenomics assembly via succinct de Bruijn graph. Bioinformatics, 31(10), 1674–1676.

Li, H., & Durbin, R. (2009). Fast and accurate short read alignment with Burrows–Wheeler transform. Bioinformatics, 25(14), 1754–1760. 10.1093/bioinformatics/btp324

Lima-Mendez, G., Faust, K., Henry, N., Decelle, J., Colin, S., Carcillo, F., Chaffron, S., Ignacio-Espinosa, J. C., Roux, S., Vincent, F., Bittner, L., Darzi, Y., Wang, J., Audic, S., Berline, L., Bontempi, G., Cabello, A. M., Coppola, L., Cornejo-Castillo, F. M., … Raes, J. (2015). Determinants of community structure in the global plankton interactome. Science, 348(6237), 1262073. 10.1126/science.1262073

Martin, M. (2011). Cutadapt removes adapter sequences from high-throughput sequencing reads. EMBnet.journal, 17(1), 10–12.

Massana, R., Murray, A. E., Preston, C. M., & DeLong, E. F. (1997). Vertical distribution and phylogenetic characterization of marine planktonic Archaea in the Santa Barbara Channel. Applied and Environmental Microbiology, 63(1), 50–56. 10.1128/aem.63.1.50-56.1997

Milanese, A., Mende, D. R., Paoli, L., Salazar, G., Ruscheweyh, H.-J., Cuenca, M., Hingamp, P., Alves, R., Costea, P. I., Coelho, L. P., Schmidt, T. S. B., Almeida, A., Mitchell, A. L., Finn, R. D., Huerta-Cepas, J., Bork, P., Zeller, G., & Sunagawa, S. (2019). Microbial abundance, activity and population genomic profiling with mOTUs2. Nature Communications, 10(1), 1014. 10.1038/s41467-019-08844-4

Mirzaei, S., & Tefagh, M. (2024). GEM-based computational modeling for exploring metabolic interactions in a microbial community. PLOS Computational Biology, 20(6), e1012233. 10.1371/journal.pcbi.1012233

Moran, M. A., Kujawinski, E. B., Schroer, W. F., Amin, S. A., Bates, N. R., Bertrand, E. M., Braakman, R., Brown, C. T., Covert, M. W., Doney, S. C., Dyhrman, S. T., Edison, A. S., Eren, A. M., Levine, N. M., Li, L., Ross, A. C., Saito, M. A., Santoro, A. E., Segrè, D., … Vardi, A. (2022). Microbial metabolites in the marine carbon cycle. Nature Microbiology, 7(4), 508–523. 10.1038/s41564-022-01090-3

Nunes, S., Latasa, M., Gasol, J., & Estrada, M. (2018). Seasonal and interannual variability of phytoplankton community structure in a Mediterranean coastal site. Marine Ecology Progress Series, 592, 57–75. 10.3354/meps12493

Olm, M. R., Brown, C. T., Brooks, B., & Banfield, J. F. (2017). dRep: A tool for fast and accurate genomic comparisons that enables improved genome recovery from metagenomes through de-replication. The ISME Journal, 11(12), 2864–2868. 10.1038/ismej.2017.126

Oña, L., & Kost, C. (2022). Cooperation increases robustness to ecological disturbance in microbial cross-feeding networks. Ecology Letters, 25(6), 1410–1420. 10.1111/ele.14006

Pachiadaki, M. G., Sintes, E., Bergauer, K., Brown, J. M., Record, N. R., Swan, B. K., Mathyer, M. E., Hallam, S. J., Lopez-Garcia, P., Takaki, Y., Nunoura, T., Woyke, T., Herndl, G. J., & Stepanauskas, R. (2017). Major role of nitrite-oxidizing bacteria in dark ocean carbon fixation. Science, 358(6366), 1046–1051. 10.1126/science.aan8260

Paerl, R. W., Curtis, N. P., Bittner, M. J., Cohn, M. R., Gifford, S. M., Bannon, C. C., Rowland, E., & Bertrand, E. M. (2023). Use and detection of a vitamin B1 degradation product yields new views of the marine B1 cycle and plankton metabolite exchange. mBio, e00061–23. 10.1128/mbio.00061-23

Paerl, R. W., Sundh, J., Tan, D., Svenningsen, S. L., Hylander, S., Pinhassi, J., Andersson, A. F., & Riemann, L. (2018). Prevalent reliance of bacterioplankton on exogenous vitamin B1 and precursor availability. Proceedings of the National Academy of Sciences, 115(44). 10.1073/pnas.1806425115

Panzeca, C., Beck, A. J., Tovar-Sanchez, A., Segovia-Zavala, J., Taylor, G. T., Gobler, C. J., & Sañudo-Wilhelmy, S. A. (2009). Distributions of dissolved vitamin B12 and Co in coastal and open-ocean environments. Estuarine, Coastal and Shelf Science, 85(2), 223–230. 10.1016/j.ecss.2009.08.016

Parks, D. H., Chuvochina, M., Waite, D. W., Rinke, C., Skarshewski, A., Chaumeil, P.-A., & Hugenholtz, P. (2018). A standardized bacterial taxonomy based on genome phylogeny substantially revises the tree of life. Nature Biotechnology, 36(10), 996–1004. 10.1038/nbt.4229

Qin, W., Heal, K. R., Ramdasi, R., Kobelt, J. N., Martens-Habbena, W., Bertagnolli, A. D., Amin, S. A., Walker, C. B., Urakawa, H., Könneke, M., Devol, A. H., Moffett, J. W., Armbrust, E. V., Jensen, G. J., Ingalls, A. E., & Stahl, D. A. (2017). Nitrosopumilus maritimus gen. Nov., sp. Nov., Nitrosopumilus cobalaminigenes sp. Nov., Nitrosopumilus oxyclinae sp. Nov., and Nitrosopumilus ureiphilus sp. Nov., four marine ammonia-oxidizing archaea of the phylum Thaumarchaeota. International Journal of Systematic and Evolutionary Microbiology, 67(12), 5067–5079. 10.1099/ijsem.0.002416

Ramond, P., Galand, P. E., & Logares, R. (2025). Microbial functional diversity and redundancy: Moving forward. FEMS Microbiology Reviews, 49, fuae031. 10.1093/femsre/fuae031

Rapala-Kozik, M. (2011). Vitamin B1 (Thiamine). En Advances in Botanical Research (Vol. 58, pp. 37–91). Elsevier. 10.1016/B978-0-12-386479-6.00004-4

Reshef, D. N., Reshef, Y. A., Finucane, H. K., Grossman, S. R., McVean, G., Turnbaugh, P. J., Lander, E. S., Mitzenmacher, M., & Sabeti, P. C. (2011). Detecting Novel Associations in Large Data Sets. Science, 334(6062), 1518–1524. 10.1126/science.1205438

Santoro, A. E., Bayer, B., Elling, F. J., & Pearson, A. (2021). Candidatus Nitrosopelagicus. En W. B. Whitman (Ed.), Bergey’s Manual of Systematics of Archaea and Bacteria (1.a ed., pp. 1-13). Wiley. 10.1002/9781118960608.gbm01969

Sañudo-Wilhelmy, S. A., Cutter, L. S., Durazo, R., Smail, E. A., Gómez-Consarnau, L., Webb, E. A., Prokopenko, M. G., Berelson, W. M., & Karl, D. M. (2012). Multiple B-vitamin depletion in large areas of the coastal ocean. Proceedings of the National Academy of Sciences, 109(35), 14041–14045. 10.1073/pnas.1208755109

Sañudo-Wilhelmy, S. A., Gómez-Consarnau, L., Suffridge, C., & Webb, E. A. (2014). The role of B vitamins in marine biogeochemistry. Annual Review of Marine Science, 6, 339–367. 10.1146/annurev-marine-120710-100912

Sathe, R. R. M., Paerl, R. W., & Hazra, A. B. (2022). Exchange of Vitamin B_1_ and Its Biosynthesis Intermediates Shapes the Composition of Synthetic Microbial Cocultures and Reveals Complexities of Nutrient Sharing. Journal of Bacteriology, 204(4), e00503–21. 10.1128/jb.00503-21

Steele, J. A., Countway, P. D., Xia, L., Vigil, P. D., Beman, J. M., Kim, D. Y., Chow, C.-E. T., Sachdeva, R., Jones, A. C., Schwalbach, M. S., Rose, J. M., Hewson, I., Patel, A., Sun, F., Caron, D. A., & Fuhrman, J. A. (2011). Marine bacterial, archaeal and protistan association networks reveal ecological linkages. The ISME Journal, 5(9), 1414–1425. 10.1038/ismej.2011.24

Steinegger, M., & Söding, J. (2017). MMseqs2 enables sensitive protein sequence searching for the analysis of massive data sets. Nature Biotechnology, 35(11), 1026–1028. 10.1038/nbt.3988

Steinegger, M., & Söding, J. (2018). Clustering huge protein sequence sets in linear time. Nature Communications, 9(1), 2542. 10.1038/s41467-018-04964-5

Suffridge, C. P., Bolaños, L. M., Bergauer, K., Worden, A. Z., Morré, J., Behrenfeld, M. J., & Giovannoni, S. J. (2020). Exploring Vitamin B1 Cycling and Its Connections to the Microbial Community in the North Atlantic Ocean. Frontiers in Marine Science, 7, 606342. 10.3389/fmars.2020.606342

Suffridge, C. P., Gómez-Consarnau, L., Monteverde, D. R., Cutter, L., Arístegui, J., Alvarez-Salgado, X. A., Gasol, J. M., & Sañudo-Wilhelmy, S. A. (2018). B Vitamins and Their Congeners as Potential Drivers of Microbial Community Composition in an Oligotrophic Marine Ecosystem. Journal of Geophysical Research: Biogeosciences, 123(9), 2890–2907. 10.1029/2018JG004554

Sultana, S., Bruns, S., Wilkes, H., Simon, M., & Wienhausen, G. (2023). Vitamin B12 is not shared by all marine prototrophic bacteria with their environment. The ISME Journal, 17(6), 836–845. 10.1038/s41396-023-01391-3

Sunagawa, S., Mende, D. R., Zeller, G., Izquierdo-Carrasco, F., Berger, S. A., Kultima, J. R., Coelho, L. P., Arumugam, M., Tap, J., Nielsen, H. B., Rasmussen, S., Brunak, S., Pedersen, O., Guarner, F., De Vos, W. M., Wang, J., Li, J., Doré, J., Ehrlich, S. D., … Bork, P. (2013). Metagenomic species profiling using universal phylogenetic marker genes. Nature Methods, 10(12), 1196–1199. 10.1038/nmeth.2693

Uritskiy, G. V., DiRuggiero, J., & Taylor, J. (2018). MetaWRAP—a flexible pipeline for genome-resolved metagenomic data analysis. Microbiome, 6(1), 158. 10.1186/s40168-018-0541-1

Vallina, S. M., Gaborit, C., Marrase, C., Gasol, J. M., Bahamon, N., Follows, M. J., Le Gland, G., & Cermeño, P. (2023). Seasonal dynamics of phytoplankton community assembly at the Blanes Bay Microbial Observatory (BBMO), NW Mediterranean Sea. Progress in Oceanography, 219, 103125. 10.1016/j.pocean.2023.103125

Wienhausen, G., Moraru, C., Bruns, S., Tran, D. Q., Sultana, S., Wilkes, H., Dlugosch, L., Azam, F., & Simon, M. (2024). Ligand cross-feeding resolves bacterial vitamin B12 auxotrophies. Nature, 629(8013), 886–892. 10.1038/s41586-024-07396-y

Wu, Y.-W., Simmons, B. A., & Singer, S. W. (2016). MaxBin 2.0: An automated binning algorithm to recover genomes from multiple metagenomic datasets. Bioinformatics, 32(4), 605–607. 10.1093/bioinformatics/btv638

Zehr, J. P., & Kudela, R. M. (2011). Nitrogen Cycle of the Open Ocean: From Genes to Ecosystems. Annual Review of Marine Science, 3(1), 197–225. 10.1146/annurev-marine-120709-142819

Zhou, Z., Tran, P. Q., Cowley, E. S., Trembath-Reichert, E., & Anantharaman, K. (2025). Diversity and ecology of microbial sulfur metabolism. Nature Reviews Microbiology, 23(2), 122–140. 10.1038/s41579-024-01104-3

Zoccarato, L., Sher, D., Miki, T., Segrè, D., & Grossart, H.-P. (2022). A comparative whole-genome approach identifies bacterial traits for marine microbial interactions. Communications Biology, 5(1), 276. 10.1038/s42003-022-03184-4

