## Supplementary Figure for "Metagenomic-based network analysis reveals the importance of vitamin cross-feeding in marine microbial assemblages"

**Supplementary information**

**Supplementary Table 1.** Edge density between MAGs with varying completeness of cobalamin and thiamine biosynthesis pathways. For cobalamin, partial synthesizers include MAGs capable of forming the corrin ring and assembling the nucleotide loop, but lacking the ability to activate the lower ligand. For thiamine, pyrimidine, and thiazole auxotrophs lack the ability to synthesize the pyrimidine (HMP) and thiazole (THZ) vitamers, respectively. Dual auxotrophs cannot synthesize either vitamer but retain the ability to combine both into thiamine. Complete auxotrophs lack any genetic capacity for thiamine biosynthesis.

| **Cobalamin** | | | | |
| --- | --- | --- | --- | --- |
| ***Edge Density*** | Complete Synthesizers | Partial Synthesizers | Lower Ligand Activators |  |
| Complete Synthesizers |  |  |  |  |
| Partial Synthesizers | 0.035 |  |  |  |
| Lower Ligand  Activators | 0.038 | 0.049 |  |  |
| Auxotrophs | 0.032 | 0.015 | 0.022 |  |
| **Thiamine** | | | | |
| ***Edge Density*** | Complete Synthesizers | Pyrimidine Auxotrophs | Thiazole Auxotrophs | Dual Auxotrophs |
| Complete Synthesizers |  |  |  |  |
| Pyrimidine Auxotrophs | 0.085 |  |  |  |
| Thiazole Auxotrophs | 0.020 | 0.043 |  |  |
| Dual Auxotrophs | 0.089 | 0.082 | 0.032 |  |
| Complete Auxotrophs | 0.075 | 0.053 | 0.019 | 0.072 |

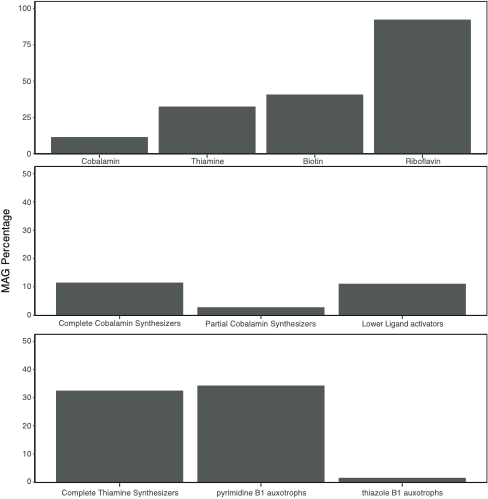

**Supplementary Figure 1.** Percentages of MAGs with different vitamin synthesis pathways. (Top) percentage of MAGs over the entire MAG pull with complete pathways for Cobalamin, Thiamine, Biotin, and Riboflavin biosynthesis. (Middle) Percentage of MAGs with complete and partial cobalamin synthesis pathways and percentage of lower ligand activators. (Down) Percentage of MAGs with a complete Thiamine biosynthesis pathway and percentage of MAGs classified as pyrimidine (HMP) and thiazole (THZ) auxotrophs.

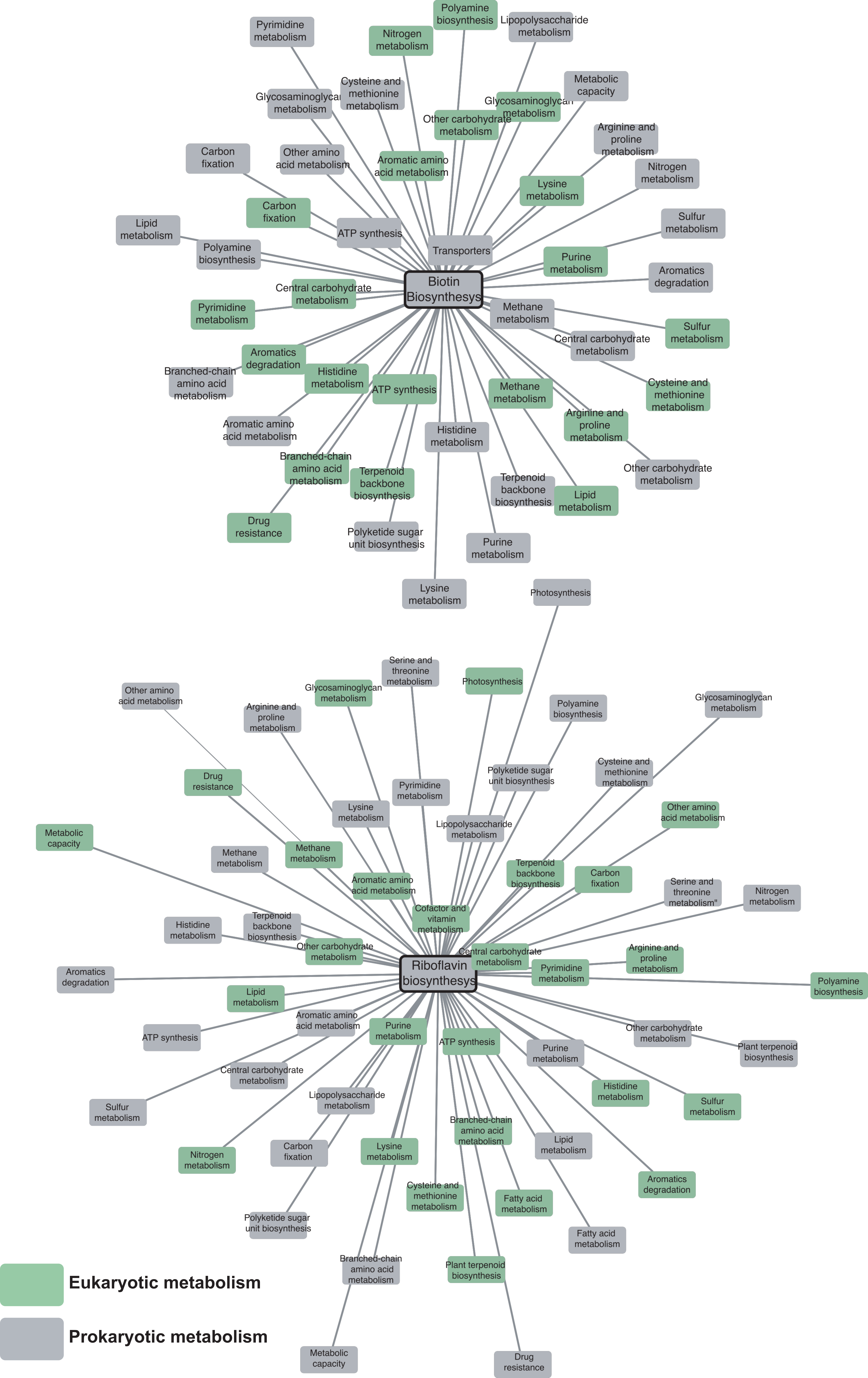

**Supplementary figure 2.** Co-occurrence networks generated with MINE, based on distinct KOs involved in the biosynthesis of biotin (top) and riboflavin (bottom). All known biosynthetic pathways for each vitamin were considered for KO selection. Vitamin synthesis KOs are connected to various prokaryotic (grey) and eukaryotic (green) metabolic pathways. Node proximity reflects the number of edges per node, indicating the connectivity within each metabolic context.

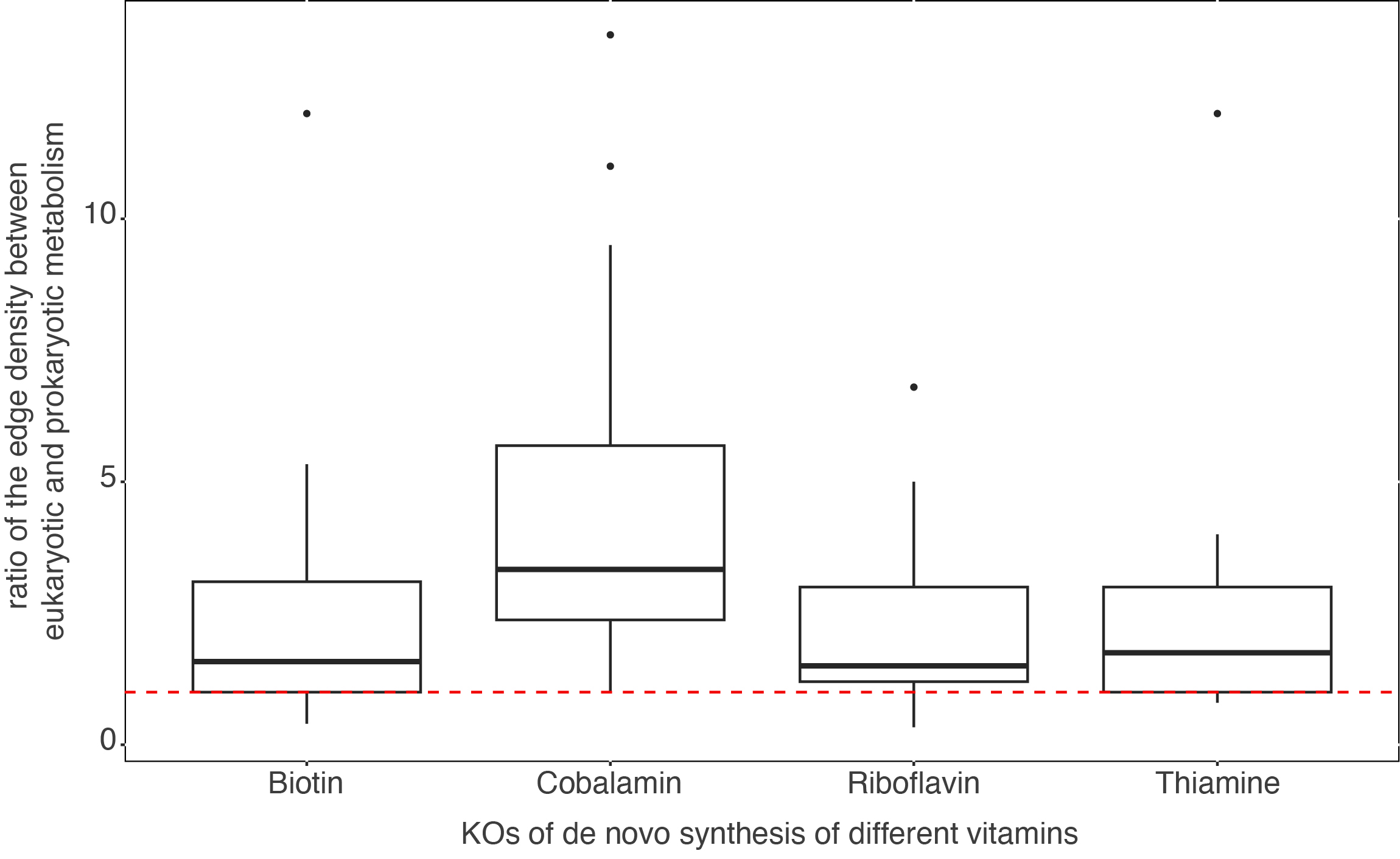

**Supplementary Figure 3.** Boxplot showing the **ratio of edge density** between **eukaryotic and prokaryotic metabolism** associated with the **prokaryotic synthesis of the four main vitamins**. A higher ratio indicates a greater edge density between **vitamin biosynthesis genes** and **eukaryotic metabolic pathways**.

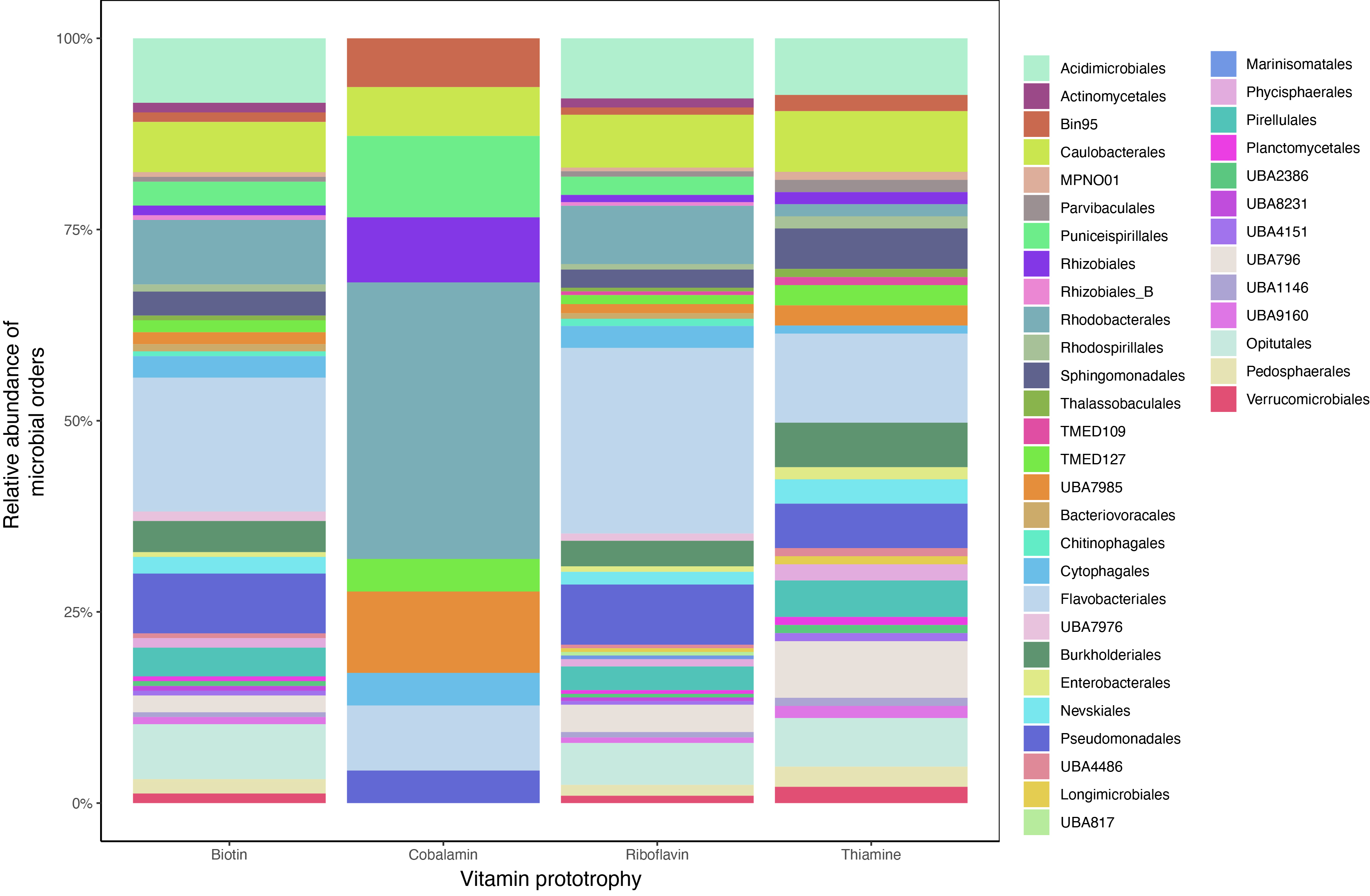

**Supplementary Figure 4.** Relative abundances of MAGs (**RPKG**) from different microbial orders with the ability to **completely or partially synthesize** the four main vitamins.

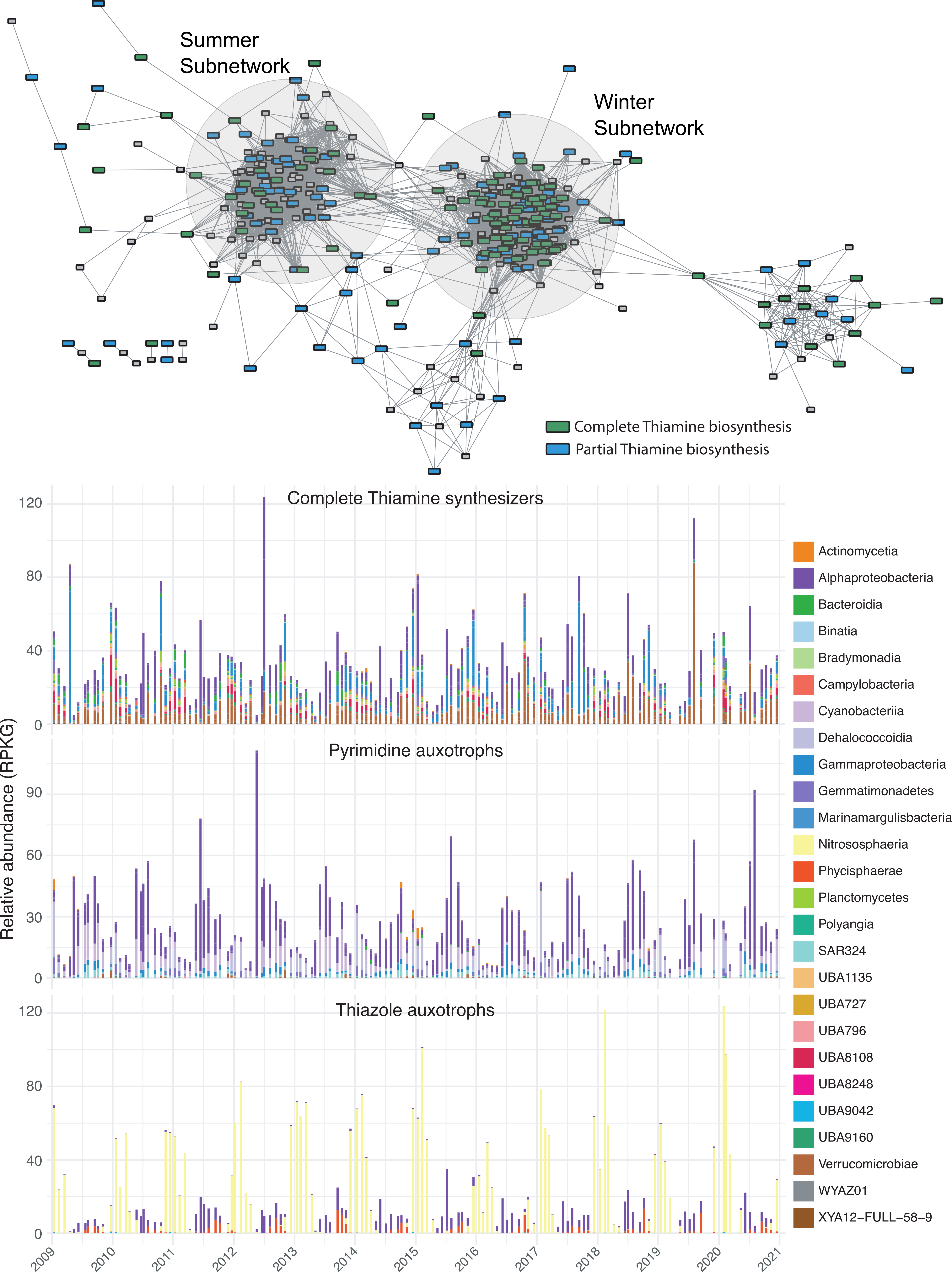

**Supplementary figure 5.** (Top) Association network of complete MAGs (>90% completeness, <10% Contamination), categorized by thiamin biosynthesis completeness: **complete** (green) and **partial** (blue). (Bottom) Relative abundance of each MAG based on their biosynthesis completeness: Complete synthesizers, pyrimidine auxotrophs, and thiazole auxotrophs.
